# Molecular relationships between SARS-CoV-2 Spike protein and LIFR, a pneumonia protective IL-6 family cytokine receptor

**DOI:** 10.1101/2021.04.18.440296

**Authors:** Mtakai Ngara, Geoffrey Henry Siwo

## Abstract

The fine-tuned control of immune responses is attained by pairs of activating and inhibitory signaling receptors which modulate the quality and magnitude of immune responses. Some viruses exploit these pathways to enter host cells as well as interfere with immune responses. Here, we report that the SARS-CoV-1/2 Spike proteins (S) contain a potential inhibitory tyrosine-based immunoreceptor motif (ITIM) with the D614G variant occurring within this motif. Through an *in silico* screen of ITIM-containing human proteins, we find that the S-located ITIM is closely related to a previously reported ITIM in the cytoplasmic tail of the human Leukemia Inhibitory Factor Receptor (LIFR), a pneumonia protective IL-6 family cytokine receptor. To infer potential functional interactions between SARS-CoV-2 infection and LIFR expression, we performed single-cell transcriptome analysis of public datasets of lung tissues from healthy individuals and COVID-19 patients. We show that transcripts of *LIFR* and its ligand *LIF* are highly expressed in SARS-CoV-2 susceptible lung cells from mild and severe COVID-19 patients but not in healthy individuals. In addition, the human endogenous retroviral envelope gene (*ERVW-1*) encoding a fusogenic protein of the same functional class as the S protein, is induced in SARS-CoV-2 susceptible cell subpopulations in COVID-19 patients with no detectable expression in healthy individuals. We also report that pulmonary epithelial cells express transcripts of several immunoreceptors including the ITIM-containing antibody receptor *FCGR2B* which is detectable in healthy and severe COVID-19 cases but not in mild cases. These results suggest that molecular dysregulation of ITIM-mediated inhibitory signaling by the SARS-CoV-2 S protein may play a role in COVID-19 immunopathology.

## Introduction

Coronavirus disease 19 (COVID-19) patients exhibit varying degrees of immune system dysfunction including potential cytokine storms^1^, acute respiratory distress syndrome (ARDS)^2^ and the production of autoantibodies against several self-antigens^3–5^. This implicates dysregulation of immune homeostasis and self-tolerance in the pathology of the disease, an area that remains understudied. Central to immune homeostasis is the coordination between activating and inhibitory signaling which ensures proportionate responses to the damage posed by pathogens during infection while maintaining tolerance to self-antigens. In immune cells this is attained by paired cell surface receptors that are either activating or inhibitory in function. Activating receptors recruit adaptor proteins containing the immunoreceptor tyrosine-based activation motifs (ITAMs) or in some cases contain ITAMs in their own cytoplasmic tails^6^. The ligation and clustering of activating receptors leads to the phosphorylation of ITAMs thereby creating docking sites for SH2-domain containing kinases which subsequently phosphorylate downstream substrates and eventually trigger effector functions such as phagocytosis and production of cytokines^6^. In contrast to activating receptors, inhibitory receptors contain immunoreceptor tyrosine-based inhibitory motifs (ITIM)^7^. Ligation of inhibitory receptors leads to phosphorylation of ITIMs in their cytoplasmic tails followed by the recruitment of SH2-domain containing cytoplasmic phosphatases (SHP2). This process eventually results in the dephosphorylation of downstream effectors and downregulation of immune responses or maintenance of self-tolerance^7^. While ITAMs are typically involved in immune activation they can also be inhibitory in some cases^8^. Certain pathogens including viruses and bacteria can exploit inhibitory signaling, for example, by mimicking ITIM/ITAM sequences in host regulatory receptors to dysregulate immune responses ^9,10^.

We hypothesized that the SARS-CoV-1/2 S proteins could potentially modulate immune signaling pathways through highly conserved short amino sequences that mediate interactions between human proteins, commonly referred to as short linear motifs (SLiMs)^11–13^. Because of their short nature (3-10 amino acids), SLiMs can rapidly emerge in viral genomes through convergent evolution and enable viral proteins to physically interact with host proteins^14^. Since the S protein is critical for cell entry by coronaviruses, elicits strong neutralizing antibodies, and has been associated with antibody-mediated enhancement (ADE) in SARS-CoV-1^15,16^, an understanding of its potential role, if any, in subversion of host immune responses is needed. We investigated whether the SARS-CoV-1/2 S proteins contain specific sequence motifs that could mimic ITIM and ITAM motifs. We focused on these two motifs because: i) They are central to balancing activating and inhibitory signals which is crucial to maintenance of self-tolerance and without which autoimmunity occurs^6^; ii) The ITAM-containing receptor (FCGR2A) and ITIM-containing (LILRB1) facilitate dengue virus ADE through binding to antibody F_c_ and by enabling endosomal escape of the virus, respectively^17–20^; iii) Some viruses mimic these motifs to downregulate immune responses^9^ and, iv) Exogenously delivered synthetic peptides containing ITIM motifs interfere with immune signaling demonstrating that standalone peptides containing ITIM motifs can effectively disrupt inhibitory signaling^21,22^.

## Results

### Spike protein contains a potential human immunoreceptor tyrosine-based inhibitory (ITIM) motif with high similarity to an ITIM sequence in LIFR

We screened SARS-CoV-1/2 S proteins against the Eukaryotic Linear Motif (ELM) database^23,24^ to identify potential SLiMs that may mediate interactions with components of the immune system. A number of SLiMs associated with interactions involving immune signaling pathways were detected in several host proteins including STAT3, STAT5, TRAF-2 binding motifs, TRAF6 binding site for Tumor Necrosis Factor 6 and ITIM (Supplementary Table 1). The vast majority of the identified SLiMs are not directly involved in immune signaling and include the recently identified RGD motif in SARS-CoV-2 S protein that binds to host integrins^25^. Whether some of these identified motifs in the virus are functional will need experimental validation. Here, we focus on the ITIM-like sequence (610-VLYQDV-615, coordinates relative to SARS-CoV-2 S; *P* = 3e-04) for the following reasons. First, antibodies to SARS-CoV-1 S derived peptide (611-LYQDVN-616) that overlaps the predicted ITIM were previously associated with ADE both in *in vitro* experiments and *in vivo* studies in non-human primate^15^. Second, the ITIM containing receptor LILRB1 has been reported to facilitate escape of dengue virus from lysosomal degradation and inhibit induction of interferon stimulated genes (ISGs) during ADE mediated viral entry into cells via the ITAM-containing FCGR2A receptor^19,20^. While ADE has not been observed in SARS-CoV-2 infections or vaccine recipients, the dysregulation of inhibitory signaling could lead to immunopathological reactions such as autoantibody production and cytokine storms that have been reported in COVID-19^1,5^. Third, the D614G mutation^26–28^ in SARS-CoV-2 lies within the predicted ITIM motif. Since ITIMs are crucial for inhibitory signaling of immune responses, mimicry by a pathogen could interfere with this process, leading to uncontrolled immune responses. Indeed, a few bacterial and viral proteins containing ITIM sequences have been established to use the motifs to interfere with host immune signaling pathways^9,10,29^.

Next, we asked whether the SARS-CoV-1/2 ITIM motif shares high sequence similarity to ITIM-like sequences located in the cytoplasmic tails of specific human proteins identified by previous computational studies^30^. Out of 109 proteins in the human proteome containing putative ITIM motifs^30^, the LIFR ITIM (VIYIDV) is highly similar to the wild-type SARS-CoV-1/2 ITIM sequence (VLYQDV), with only 2 amino differences between the two. Another host protein with a closely related ITIM is the platelet derived growth factor receptor, beta precursor (PDGFRB) with the sequence VLYTAV^30^. Previous studies indicate that LIFR is required for lung protection during pneumonia, for example, in respiratory syncytial virus infections (RSV)^31–34^. Furthermore, its ligand, LIF, is one of the cytokines in the IL-6 family and is increased in the lungs of patients with ARDS^35,36^, which has been observed in COVID-19 patients^2,37,38^. Notably, because the D614G variant is located within the SARS-CoV-2 ITIM, the mutational distance between the LIFR ITIM and the corresponding ITIM sequence in D614G viruses increases from 2 to 3 which could reduce the potential molecular mimicry or subversion of LIFR by the D614G associated ITIM. A number of studies suggest that D614G variants are highly transmissible^26,27,39,40^ and that the mutation increases infectivity to lung cells^40–46^ implying the possibility that this sequence region is under selective pressure. However, to date no associations have been found between this mutation and COVID-19 disease severity^26,28^.

### Single-cell transcriptome analysis of LIFR, LIF and SARS-CoV-2 entry receptors in the human cell atlas and lung cell transcriptome

To further explore the possibility of biological interactions between LIFR and SARS-CoV-2, we analyzed LIFR expression in single-cell transcriptomes of human tissues using data from the human cell atlas (HCA)^47^ and lung single cell transcriptome from the Madissoon et al study^48^ as well as a recently published single-cell bronchoalveolar transcriptome of COVID-19 patients and healthy controls^49^ (Fig. 1-2, Supplementary Fig. S1 and S2). In addition, we included single-cell expression levels of genes important for SARS-CoV-2 cell entry (*ACE2, TMPRSS2, NRP1, NRP2*)^50–54^, those previously implicated in ADE via Fc receptor binding and/or signaling via ITIM/ITAM sequences (*FCGR2A*^17,18^, *FCGR2B*^7^, *LILRB1*^19,20^, *NKG2A*^55^, *MyD88* and *CD300LF/IREM-1*^21,22^) or modulating ADE (Fut8)^56,57^ as well as key cytokines (IFNG and IFNB). Additionally, we included *ERVW-1* which encodes the human endogenous retrovirus W (HERV-W) envelope protein or syncytin, one of a few host proteins that are functionally similar to the viral spike protein in catalyzing fusion of cell membranes. Further details on the selected genes are described in the Methods section.

**Figure 1:**
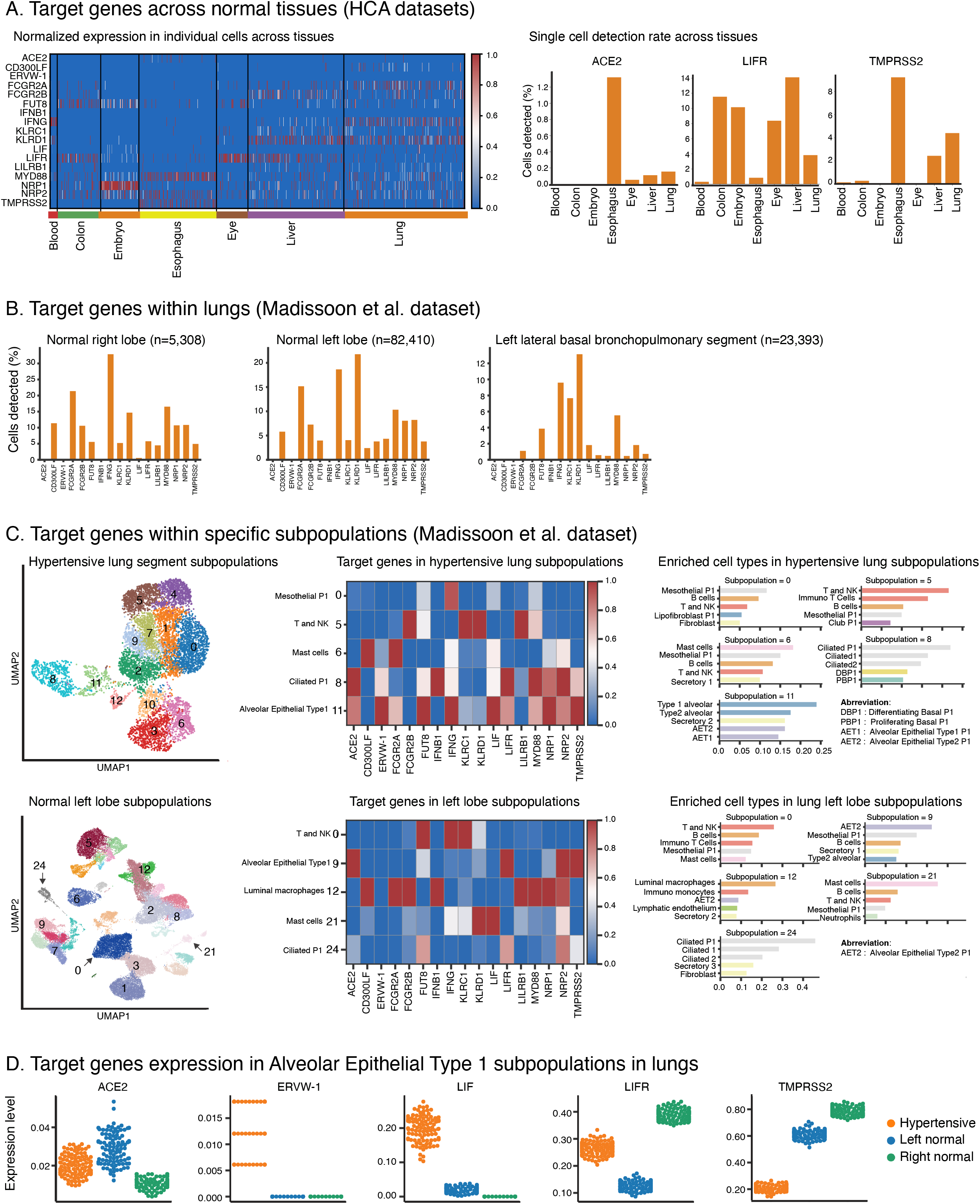
Immunopathology analysis in human cell atlas datasets. **a** Single-cell expression of target genes across normal tissues (HCA datasets). The heatmap depicts the normalized expression level of target genes (Y-axis) in single cells from distinct tissues (X-axis) in the human cell atlas (HCA) datasets. Only single cells with at least one of the target genes detected and at least 300 genes detected were included. The gene expression in each cell was transformed to standard scale. Single cell detection rates across tissues are shown in the bar plots by the percentage of single cells (Y-axis) in each tissue from the HCA datasets (X-axis) in which three of the target genes (*ACE2, LIFR* and *TMPRSS2*) are detected (counts >=1). Only single cells with at least 300 detected genes were included in the analysis. **b** Expression of target genes within lungs from Madissoon et al study^48^. Three bar plots of the percentage (Y-axis) of lung-derived single cells from the HCA datasets (X-axis) in which the candidate genes (X-axis) are detected (counts >=1). The cells were derived from the right lobe (left panel), left lobe (middle panel) and from a suspected hypertensive lung from the left lateral basal bronchopulmonary segment (right panel). Above each plot are the total number of single cells analyzed which were cells with at least 300 genes detected. **c**. Single-cell expression of the candidate genes within specific subpopulations in the left lobe of normal lung and suspected hypertensive lung. Distinct cell subpopulation in UMAP plots are labeled by numbers. Only cells with 1000 or more genes detected (counts>=1) with mitochondrial fraction<=0.2 fraction were included. The counts per cells were normalized to 10,000 then dimensions reduced to 20 principal components (PCA) and projected on two UMAP dimensions based on significantly variable genes. The neighborhood graphs were based on UMAP embedding and Euclidean distance with clustering using Louvain-graph algorithm. The heatmaps show the mean expression (on standard scale) of the candidate genes from a subset of subpopulations from the left lobe. Each subpopulation is labelled by the enriched cell type as described in the Methods section. The bar plots show five of the top enriched cell types in a subset of subpopulations from the left lobe of the lung. The score on the X-axis of the bar plots represents the overlap between top 100 ranked genes per subpopulation against the reference markers for each cell type. For the normal left lobe (upper panel), the number of cells per subpopulation are: 0 (n= 3995), 1 (n= 3573), 2 (n= 3499),3 (n= 2837),4 (n= 2702),5 (n= 2684),6 (n= 1765),7 (n= 1487),8 (n= 1363),9 (n= 1250),10 (n= 1225),11 (n= 1188),12 (n= 1183),13 (n= 1081),14 (n= 1036),15 (n= 972),16 (n= 958),17 (n= 856),18 (n= 829),19 (n= 653),20 (n= 634),21 (n= 619),22 (n= 556),23 (n= 477),24 (n= 438),25 (n= 321),26 (n= 103) and 27 (n= 70). For the hypertensive lung (upper panel), the number of cells per subpopulation are: 0(n=1268), 1 (n=914), 2 (n=807), 3 (n=807), 4 (n=801), 5 (n=744), 6 (n=737), 7(n=550), 8(n=517), 9(n=453), 10(n=264), 11(n=253) and 12(n=128). **d**. Target genes expression in AET1 subpopulation. Strip plot showing normalized expression (Y-axis) of five candidate genes (*ERVW-1, LIF, TMPRSS2, LIFR* and *ACE2*) across subpopulations enriched for AET1 cells. The points are colored based on the lung section from which the subpopulation is derived i.e. left hypertensive lung(orange), left normal lobe (blue) and right normal lobe(green).

The overall expression of the selected target genes across human tissues shows that the key SARS-CoV-2 entry factors (ACE2 and TMPRSS2) are sparsely expressed in normal tissues with the highest detectable single-cell expression of ACE2 in the esophagus (∼1.4% of cells) followed by lungs (∼0.18% of cells), liver (∼0.1% of cells) and eye (∼0.05% of cells) (Fig. 1 A, HCA data). As lungs are the primary tissue targeted by SARS-CoV-2, we also analyzed the expression of the selected candidate genes in the Madissoon et al study^48^ (Fig. 1B). The percentage of cells expressing *ACE2, TMPRSS2* and *LIFR* in lung tissue was highly comparable between the HCA and the Madissoon et al study datasets with extremely low ACE2 expression in both. This is consistent with previously published studies^58,59^. Next, we applied unsupervised approach based on uniform manifold learning (UMAP)^60^ with Louvain algorithm for community detection^61^, to the previously published lung single-cell transcriptome by Madissoon et al^48^. We established 27 and 12 cell subpopulations based on data from the normal left lobe of the lungs and a suspected hypertensive left lung, respectively (Fig. 1 C). To determine whether these subpopulations are enriched with specific cell types, we obtained cell types markers as described in UCSC cell browser (https://cells.ucsc.edu/). We then computed an overlap score between the top 100 markers of each subpopulation and each cell type’s markers using Scanpy package^62^. We identified enrichments of cell types in each of the cell subpopulations including enrichments in Alveolar Type 1/ 2 (AET1/2) and Ciliated cells that are SARS-CoV-2 infection primary targets. The expression of the candidate genes in 5 subpopulations enriched with distinct cell types is shown for normal lungs by the heatmap and bar plots in Fig. 1C and a suspected hypertensive lung in Fig. 1D (See Supplementary Fig. S3 for detailed data on all cell type enrichments across all the cell subpopulations detected).

SARS-CoV-2 primarily infects lung epithelial cells here represented by AET1/2 and ciliated cells (ciliated P1/P2, ciliated 1/2). Consistent with this, the main entry receptor of the virus ACE2 and the TMPRSS2 protease required to activate the fusogenic activity of the S protein are both expressed in AET1/2 and ciliated cells (Fig. 1C and Supplementary Fig. S3). In addition, these cells expressed the recently identified SARS-CoV-2 entry receptor, NRP1, as well as its close relative NRP2^53,54^ (Fig. 1C and Supplementary Fig. S3). Overall, AET1 and ciliated P1 cells have distinct gene expression profiles of the candidate genes in normal vs hypertensive lung (heatmaps in Fig. 1C). Surprisingly, AET1 and ciliated P1-the main cell subpopulations expressing both *ACE2* and *TMPRSS2* and therefore susceptible to SARS-CoV-2 -also show enhanced expression of *ERVW-1, LIF, LIFR, FUT8, IFNB1, IFNG, KLRC, MYD88* in the hypertensive lungs compared to normal lungs. In contrast, *TMPRSS2, KLRD1* and *NRP2* are downregulated in AET1, and the remaining genes including ACE2 are comparably expressed in normal vs hypertensive lungs (Fig. 1D; detailed in Supplementary Fig. S3 and S4). Notably, the hypertensive lung was originally identified as healthy in the Madissoon et al study^48^. Interestingly, the expression levels of the candidate genes from the right and left normal lungs are distinct potentially due to differences in lung volume, implying that gene expression analyses of lung cells need to take this into account. Out of the genes analyzed, *ERVW-1* has a unique expression pattern -it is not expressed at all in both left and right normal lungs and is weakly but detectably expressed in the hypertensive lung, and even so only in AET1 and ciliated P1 cells (Fig. 1D).

### Single-cell transcriptome analysis of LIFR, LIF and SARS-CoV-2 entry receptors in the lungs of COVID-19 patients

The expression of a subset of the candidate genes in host cells of SARS-CoV-2 increases the likelihood of biologically relevant interactions with the viral spike protein. Therefore, we next examined the expression of the candidate genes in an independent single-cell RNAseq data from the bronchoalveolar lavage fluid (BALF) of three cases of mild COVID-19, six severe cases and three healthy controls using a recently published dataset^49^ (QC analysis shown in Supplementary Fig. S5). Clustering of the single cell transcriptomes resulted in distinct groupings with a clear separation of cells from the healthy controls and a closer relationship between the mild and severe cases (Fig. 2A; Supplementary Fig. 6 A). In addition to this separation, single cells from individual subjects within each group are also separated with cells from the same individual generally closer to each other (Fig. 2A). As expected, high levels of SARS-CoV-2 transcripts are detectable in the single cells of severe cases with relatively fewer transcripts in the mild cases and none in the healthy controls (Fig. 2A; Supplementary Fig. S6 B-C). A summary view of the expression of each of the candidate genes in single cells from each group is shown in the track plots and UMAP plots in Fig. 2B. It is evident that *ACE2* expression is especially higher in severe COVID-19 cases relative to mild disease with no detectable expression in the lung from the healthy cases (Fig. 2B; Supplementary Fig. 6C). *TMPRSS2* on the other hand shows considerable expression in the lung tissue of severe and mild cases with a low expression in the healthy cases (Fig. 2B; heatmaps Fig. 2C; Supplementary Fig. 6C). We also observed that expression of *ERVW-1* and *IFNB1* is only detectable in severe cases (Fig. 2B track plots and heatmaps Fig. 2C; Supplementary Fig. 6C) which is noteworthy considering that *ERVW-1* expression induces double-stranded RNA antiviral response^63–65^. Furthermore, *IFNB1* transcript expression was recently associated with prolonged ICU stay and multiorgan involvement which are hallmarks of critical COVID-19 cases^66^. On the other hand, we found that *LIFR* expression is high in both mild and severe cases with no detectable levels in healthy cases (Fig. 2B track plots and 2C heatmaps; Supplementary Fig. 6C). In contrast, LIF is expressed in all cases even though expression in mild and severe cases is more widespread (Fig. 2B track plots and UMAP; heatmaps Fig. 2C; Supplementary Fig. 6C).

**Figure 2:**
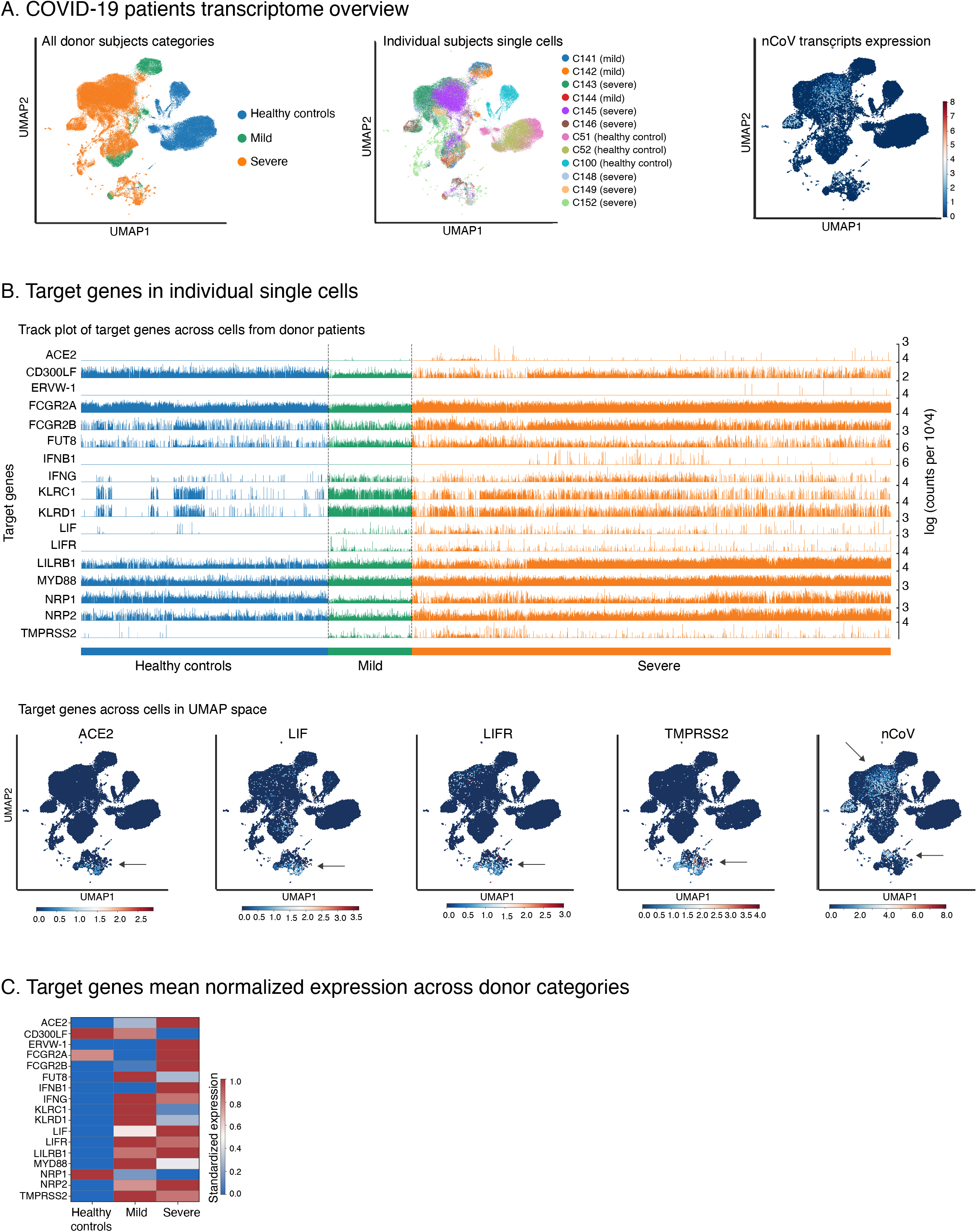
Immunopathology analysis in COVID-19 BALF datasets (Liao et al. study^49^, GSE145926). **a**. COVID-19 patients transcriptome overview based on Liao et al. study lab dataset. Top left panel-UMAP projection of the single cell transcriptomes from bronchoalveolar lavage fluid (BALF) of mild (green) and severe(orange) COVID-19 patients and healthy donors(blue). Only cells with 1000 or more genes detected (counts>=1) and reads fraction<=0.2 mapped towards mitochondrial genes. The counts per cells were normalized to 10,000 then dimensions reduced to 20 principal components (PCA analysis) and projected on two UMAP dimensions based on significantly variable genes. Middle panel-UMAP projection of the single cell transcriptomes from BALF of individual subjects from the Liao et al. study^49^. Each cell (points) is colored according to the individual subject donor with the disease status indicated in brackets. Right panel-UMAP projection of the single cell transcriptomes from BALF of individual subjects from the Liao, M. et al. study. Each cell (points) is colored according to the number of reads mapped towards the SARS-CoV2 virus ranging from low (deep blue) to high(red) expression. **b**. Target genes in individual single cells. Track plots showing the log-normalized expression (Y-axis, right margin) of the 17 target genes (Y-axis, left margin) in individual cells (X-axis) from the Liao, M. et al. study BALF datasets. Each track is colored based on the donor subject disease state with healthy controls(blue), mild (green) and severe(orange) COVID-19 patients. **c**. Target genes across cells in UMAP space. The normalized expression of 4 target genes (*ACE2, LIF, LIFR* and *TMPRSS2*) and the SARS-CoV-2 transcripts (above each panel) in the UMAP projected single cell transcriptomes from the Liao, M. et al. study BALF datasets. Each cell (points) is colored according to the expression magnitude of the target gene ranging from low (deep blue) to high (red) expression. The arrow indicates the cells subset where each target gene is highly expressed. **d**. Target genes mean normalized expression across donor categories. A heatmap showing the mean expression (on standard scale) of 17 target genes in donor subjects from the three categories namely healthy, mild and severe from the Liao et al. study COVID-19 BALF datasets.

To further examine the single cell expression levels of the candidate genes in COVID-19 infected vs. healthy lungs, we performed an analysis of cell subpopulations to determine the expression levels of the candidate genes in enriched cell types within each subpopulation (Fig. 3; Supplementary Fig. S9 for enriched cell types in each subpopulation). *ACE2* and *TMPRSS2* expression levels are extremely low in single cells from healthy individuals while relatively higher expression is observed in ciliated P1, goblet cells and alveolar type 1/2 cells of mild and severe COVID-19 cases (Fig. 3 and Supplementary Fig. 7-8). A similar pattern of single cell expression is observed for *ERVW-1, LIF* and *LIFR* (Fig. 3 and Supplementary Fig. 7-8). In contrast, while *FCGR2A* and *FCGR2B* were detectable across various cell types in both healthy and COVID-19 cases, the single cell expression of these genes was much lower in Ciliated P1 cells of the COVID-19 mild and severe cases (Supplementary Figure S8). The expression of these antibody receptors in pulmonary epithelial cells is unexpected as most of their functions are associated with professional immune cells. The expression of the SARS-COV-2 entry receptor and protease in the same cell types as *ERVW-1, LIF* and *LIFR* increases the likelihood of interactions of the corresponding proteins. Furthermore, given that expression level of these genes is extremely low in healthy cases especially in cell types susceptible to SARS-CoV-2, there appears to be a potential link to SARS-CoV-2 infection.

**Figure 3:**
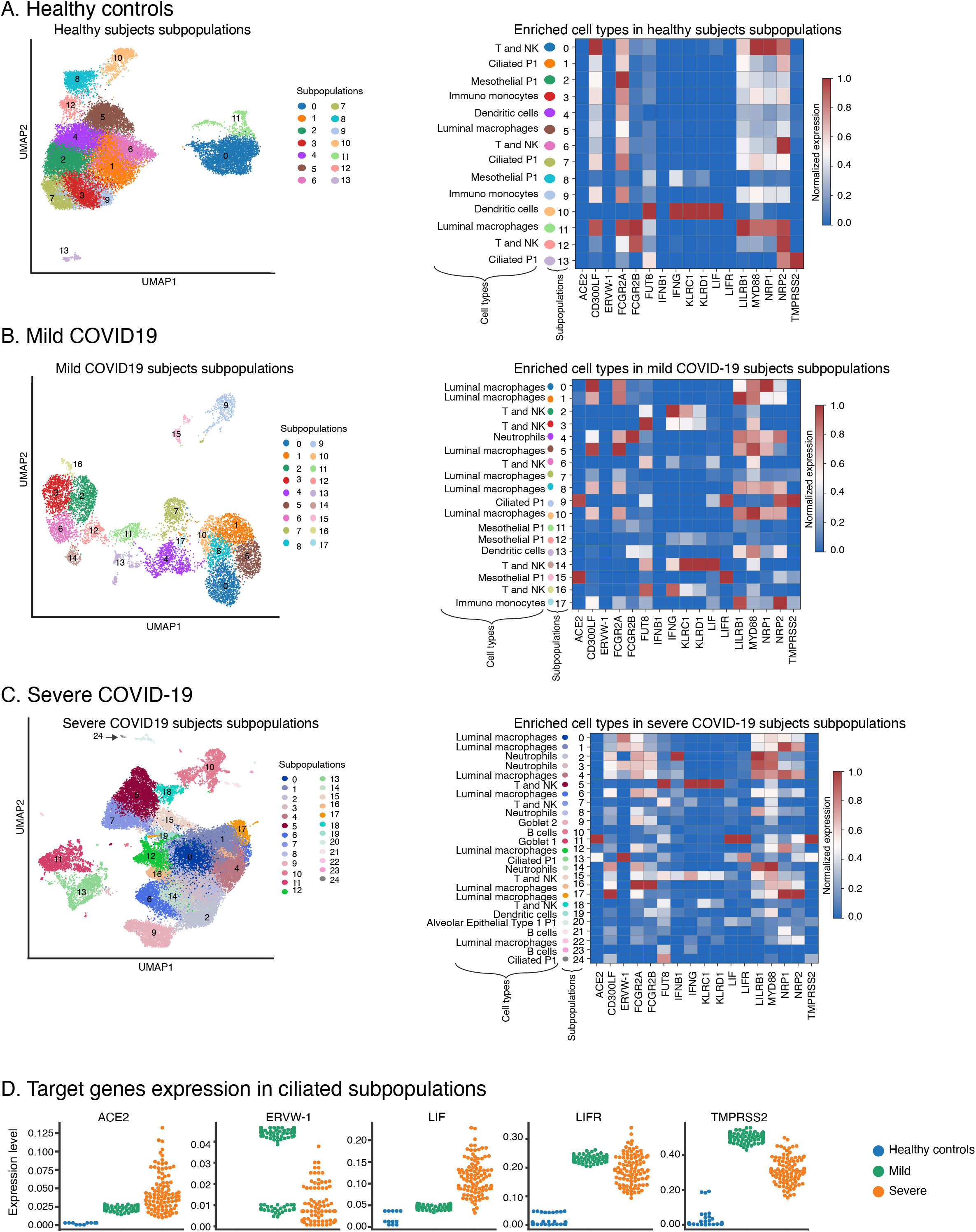
Target genes within subpopulations of COVID-19 BALF datasets (Liao et al. study^49^, GSE145926). **a**. Healthy subject subpopulations. UMAP projection of the single cell transcriptomes from BALF of healthy subjects. Each cell (points) is colored according to the subpopulation with number labels. Only cells with 1000 or more genes detected (counts>=1) with mitochondrial fraction<=0.2 fraction were included. The counts per cells were normalized to 10,000 then dimensions reduced to 20 principal components and projected on two UMAP dimensions based on significantly variable genes. The neighborhood graphs were based on UMAP embedding and Euclidean distance with clustering using Louvain-graph algorithm. The number of cells per subpopulation are: 0 (n= 3596), 1 (n= 3584), 2 (n= 3202), 3 (n= 2772), 4 (n= 2516), 5 (n= 2496), 6 (n= 1838), 7 (n= 1377), 8 (n= 1152), 9 (n= 943), 10 (n= 686), 11 (n= 490), 12 (n= 455) and 13 (n= 109). The heatmap shows the mean expression (on standard scale) of 17 target genes in the single cell subpopulations from BALF derived from healthy donors. Each subpopulation is labelled by the enriched cell type. **b**. Analysis of cell subpopulations in mild COVID-19 subjects. UMAP projection of the single cell transcriptomes from BALF of mild COVID-19 cases (Liao et al. study). Each cell (points) is colored according to the subpopulation with number labels. Only cells with 1000 or more genes detected (counts>=1) with mitochondrial fraction<=0.2 fraction were included. The counts per cells were normalized to 10,000 then dimensions reduced to 20 principal components (PCA) and projected on two UMAP dimensions based on significantly variable genes. The neighborhood graphs were based on UMAP embedding and Euclidean distance with clustering using Louvain-graph algorithm. The number of cells per subpopulation are: 0 (n= 1038), 1 (n= 917), 2 (n= 859), 3 (n= 803), 4 (n= 798), 5 (n= 690), 6 (n= 587), 7 (n= 543), 8 (n= 402), 9 (n= 379), 10 (n= 349), 11 (n= 296), 12 (n= 240), 13 (n= 213), 14 (n= 167), 15 (n= 127), 16 (n= 65), and 17 (n= 42). The heatmap shows the mean expression (on standard scale) of 17 target genes (X-axis) in the single cell subpopulations from BALF derived from mild COVID-19 subjects. Each subpopulation is labelled by the enriched cell type. **c**. Analysis of cell subpopulations in severe COVID-19 cases. UMAP projection of the single cell transcriptomes from BALF of severe cases (Liao, M. et al. study). Each cell (points) is colored according to the subpopulation with number labels. Only cells with 1000 or more genes detected (counts>=1) with mitochondrial fraction<=0.2 fraction were included. The counts per cells were normalized to 10,000 then dimensions reduced to 20 principal components and projected on two UMAP dimensions based on significantly variable genes. The neighborhood graphs were based on UMAP embedding and Euclidean distance with clustering using Louvain-graph algorithm. The number of cells per subpopulation are: 0 (n= 5821), 1 (n= 4991), 2 (n= 4866), 3 (n= 4453), 4 (n= 3845), 5 (n= 3352), 6 (n= 2749), 7 (n= 2364), 8 (n= 2308), 9 (n= 2243), 10 (n= 1771), 11 (n= 1697), 12 (n= 1489), 13 (n= 1466), 14 (n= 1350), 15 (n= 1142), 16 (n= 920), 17 (n= 609), 18 (n= 575), 19 (n= 428), 20 (n= 200), 21 (n= 109), 22 (n= 41), 23 (n= 25), and 24 (n= 23). The heatmap shows the mean expression (on standard scale) of 17 target genes in the single cell subpopulations from BALF of the severe cases. Each subpopulation is labelled by the enriched cell type. **d**. Target genes expression in ciliated subpopulations. Strip plot showing normalized expression of five target genes (*ACE2, ERVW-1, LIF, LIFR* and *TMPRSS2*) across subpopulations enriched for ciliated cell type. The points are colored based on the subjects from which the subpopulation is derived i.e. healthy controls(blue), mild(green) and severe(orange) COVID-19 patients.

## Discussion

There is increasing evidence that dysregulation of immune responses plays a critical role in COVID-19 through several mechanisms. For example, autoantibodies against a broad range of extracellular antigens such as cytokines (IFNs, IL6)^3,5^, chemokines (CXCL1, CXCL7)^5^ and several immunomodulatory cell surface receptors (NKG2A/C/E, CD38) are overexpressed in subsets of COVID-19 patients^67,68^. The molecular mechanisms through which infection by the SARS-CoV-2 virus could lead to a breakdown in tolerance remain unknown. Woodruff et al suggested that occurrence of autoimmunity in a subset of COVID-19 patients could be due to previously unknown pre-existing autoimmunity in the patient cohort or a consequence of immunological events in COVID-19 patients^4^. Activation of TLR7 by the single-stranded RNA of the virus could lead to de novo autoreactivity to self-antigens^4^. Given that ITIM signaling modulates TLR7 responses^69^ and several ITIM-containing receptors are involved in B-cell tolerance^70^, potential targeting of host ITIMs by a virus could lead to a breakdown in self-tolerance.

In this study, we found that the S protein contains a hexapeptide motif that has a high similarity to an ITIM in a host protein, LIFR. Furthermore, this hexapeptide contains the site for the D614G mutation in SARS-CoV-2. Interestingly, in SARS-CoV-1, antibodies to this peptide were associated with ADE in *in vitro* experiments and led to lung induced injury in non-human primates^15^. Our analysis of the similarity of the hexapeptide to ITIM sequences in 109 human proteins revealed that it is most closely related to an ITIM sequence located in the Leukemia Inhibitory Factor Receptor (LIFR), an interleukin-6 (IL-6) family cytokine receptor. Whether the ITIM sequence in the virus can interfere with signaling by LIFR or other host proteins requires further laboratory validation as the functional activity of the LIFR ITIM has not been verified experimentally. Typically, functional ITIM sequences are located within the cytoplasmic tail of immunoreceptors^6^. The ability of exogenously delivered ITIM containing peptides to inhibit TLR-mediated ITIM-dependent signaling^21,22^ opens the possibility that virus derived ITIMs can similarly interfere with inhibitory ITIM signaling. For this to occur, the viral proteins containing the ITIM sequences need to be present within cell types expressing host proteins with ITIMs in their cytoplasmic tails. Thus, we assessed the single cell expression of LIFR and other ITIM containing host proteins as a first step to determine whether SARS-CoV-2 infection can result in dysregulation of ITIM signaling.

To date there is limited data on the role if any of LIF/ LIFR in COVID-19^71^. Yet, LIF is known to be an important player in recovery from pneumonia^31^ and is elevated in the lungs of patients with ARDS^35^. Like all members of the IL-6 family of cytokines, LIF signals via STAT3 and utilizes glycoprotein 130 (gp130) as a receptor subunit^72^. LIF signaling is mediated by its binding to a high affinity receptor complex composed of a low affinity LIF binding chain (LIFR) and a high affinity converter subunit (gp130)^73^. Our results show that LIF is particularly elevated in mild COVID-19 pulmonary epithelial cells compared to severe or healthy cases while LIFR is elevated in both mild and severe cases (Figure 3; Supplementary Figure S8). Thus, it is possible that enhanced levels of LIF may lead to mild disease by compensating for potential interference of LIFR signaling by the viral spike ITIM sequence.

The classical view of the biological role of pulmonary epithelial cells is that they serve as a physical barrier to limit entry of pathogens or facilitate the removal of pathogens. Recent studies show that pulmonary epithelial cells play more complicated roles in inflammation, immunity and host defense^74^. Our results indicate that at least at the transcriptional level, pulmonary epithelial cells also express other key immunoreceptors including those containing ITAMs (FCGR2A), and ITIMs (LIFR, FCGR2B, LILRB1, NKG2A) implying that these cell types may also be involved in various immune responses. However, our study is limited by the use of transcriptional datasets which do not always reflect functional relevance. Interestingly, dexamethasone, a corticosteroid that improves survival of severely ill COVID-19 patients^75,76^ has been shown to enhance expression of *FCGR2B* both at the mRNA and protein levels^77,78^. It is notable that *FCGR2B* expression was detectable in ciliated cells of healthy as well as severe COVID-19 cases but not in those of mild COVID-19 cases (Supplementary Figure S8). It is also possible that subversion of host ITIMs not considered here could also contribute to immune dysregulation in COVID-19. Dysregulation of other ITIM-containing proteins such as PD-L1 expressed by airway epithelial cells ^79^ or PECAM1 which is co-located with FCGR2A in platelets^80^ may be especially important given the recent reports of vaccine induced thrombocytopenia in SARS-CoV-2 ChaAdOX1 vaccine recipients^81,82^.

The observation that expression of *ERVW-1* is induced in both mild and severe COVID-19 could also result in immune dysregulation and demonstrates a potential role of airway epithelial cells in pulmonary immune homeostasis given that ERV-derived dsRNA can trigger innate immune responses^63,64,83^. In relation to this, a recent study found that SARS-CoV-2 infection leads to increased expression of LINE-1, another type of retrotransposons in the human genome^84^. Since LINE-1 encoded reverse transcriptase can also reverse transcribe HERV-W transcripts^85^, this may increase the likelihood for HERV-W pseudogene formation. There are at least 13 loci in the human genome encoding full length HERV-W envelope protein (Env) with only one of these on chromosome 7 encoding a functional protein, syncytin that is key to placenta formation through fusion of cytotrophoblasts^86^. HERV-W Env expression has been associated with multiple sclerosis^87^ and type-1 diabetes^88^. Furthermore, Env is cytotoxic and induces production of autoantibodies that target it^89^. Such Env targeting autoantibodies in type-1 diabetes patients could cross-react with Env/ syncytin expressed in the lungs of COVID-19 patients. This could be one potential mechanism through which some comorbidities, for example, type-1 diabetes may be associated with severe COVID-19.

LIF and IL-6 share a signaling receptor (gp130) yet most studies to date on COVID-19 have focused only on IL-6 and a detailed assessment of the role of LIF /LIFR in the disease is lacking. IL-6 is one of the top cytokines that have been associated with COVID-19 disease severity yet clinical trials of therapeutic benefit of anti-IL-6 antibodies have yielded mixed results^90,91^. A dissection of the role of LIF and IL-6 in COVID-19 could guide targeting of the distinct signaling pathways for these IL-6 family cytokines as a means to develop therapeutics against COVID-19. Given that LIFR is required for protection of lungs in pneumonia and its ligand LIF has been associated with acute respiratory distress syndrome (ARDS), comorbidities such as obesity^92,93^, diabetes^94^, hypertension^95,96^, stroke^97^ and kidney failure^98^, it may provide insights into why these conditions are associated with severe COVID-19. As LIFR is also important in the proliferation of olfactory neural progenitors which influence neuronal regeneration of the olfactory epithelium after injury^99,100^, it could have a direct role in the loss of smell in COVID-19 patients. Thus, the results of this study have implications for understanding disease pathology in SARS-CoV-2.

## Supplementary Figures

**Figure S1:** Immunopathology analysis in human cell atlas (HCA) datasets. **a**. Track plot of target genes across cells across normal tissues in HCA at single cell resolution. Track plots showing the log-normalized expression (Y-axis, right margin) of the 17 target genes (Y-axis, left margin) in individual cells (X-axis) derived from different HCA tissues. Each track is colored based on the source tissue of the specific cell. **b**. Target genes detection rate in single cells normal across tissues. Bar plots show the percentage (Y-axis) of single cells in each tissue from the HCA datasets (X-axis) in which 17 of the target genes (as labelled above each figure panel) are detected (counts >=1). Only single cells with at least 300 detected genes were included in the analysis. **c**. Target genes detection in lung-derived cells (Madissoon et al study^48^). Bar plots show log counts (Y-axis) of the 17 target genes (X-axis) in single cells from three tissue segments derived from: left lateral basal broncho-pulmonary section of a subject with a suspected hypertensive disorder (n=8042), lower lobe of right lung of a normal lung parenchyma (n=38951) and lower lobe of a left normal lung parenchyma (n= 4070). **d**. Target genes detection rate in lung-derived cells (Madissoon et al study ^48^). Bar plots showing the detection rate (Y-axis) of the 17 target genes (X-axis) in single cells from three tissue segments derived from: left lateral basal broncho-pulmonary lug from a subject with suspected hypertensive disorder (n=8042), lower lobe of a normal right lung parenchyma (n=38951) and lower lobe of a normal left lung parenchyma (n= 4070).

**Figure S2:** Target genes normalized expression in individual cells in HCA tissues. Combined violin and jitter plots showing the log-normalized expression (Y-axis) of the 17 target genes (X-axis) in individual cells from the HCA normal tissues.

**Figure S3:** Target genes expression within the lung datasets (Madissoon et al study^48^). **a**. Transcript levels in cell subpopulations from a suspected hypertensive lung. The figure on the left shows enriched cell types (y-axis) and their respective scores (x-axis) using horizontal bar plots in the 12 single cell subpopulations derived from the suspected hypertensive lung segment. The bars are colored according to the cell types markers they represent and above each panel is the subpopulation’s label. Some of the names for the cell types are abbreviated in the legend below the figure panel. On the top right section of the figure is a UMAP projection of the 12 single cell subpopulations, in distinct color points, from the hypertensive lung segment and their corresponding numerical label. The heatmap shows the scaled expression level of the 17 target genes (x-axis) in the 12 subpopulations with their top scoring cell type marker (y-axis). **b**. Normal left lobe subpopulations. The figure on the left shows enriched cell type markers (y-axis) and their respective scores(x-axis) using horizontal bar plots in the 27 single cell subpopulations derived from normal lung left lobe subpopulations. The bars are colored according to the cell types markers they represent and above each panel is the subpopulation’s label. On the top right section of the figure is a UMAP projection of the 27 single cell subpopulations, in distinct color points, from the normal lung left and their corresponding numerical label. Below is a heatmap showing the scaled expression level of the 17 target genes (x-axis) in the 27 subpopulations with their top scoring cell type marker (y-axis).

**Figure S4:** Target genes expression within single cell lung datasets (Madissoon et al study^48^) of the right lobe of a normal lung. The figure on the left shows enriched cell types (y-axis) and their respective scores(x-axis) using horizontal bar plots in the 15 single cell subpopulations derived from normal lung right lobe subpopulations. The bars are colored according to the cell types markers they represent and above each panel is the subpopulation’s label. On the top right section of the figure is a UMAP projection of the 15 single cell subpopulations, in distinct color points, from the normal lung right lobe and their corresponding numerical label. The number of cells per subpopulation are: 0 (n= 526),1 (n= 491),2 (n= 469),3 (n= 409),4 (n= 358),5 (n= 346),6 (n= 341),7 (n= 217),8 (n= 149),9 (n= 118),10 (n= 114),11 (n= 75),12 (n= 61),13 (n= 56),14 (n= 32) and 15 (n= 25). Below is a heatmap showing the scaled expression level of the 17 target genes (x-axis) in the 15 subpopulations with their top scoring cell type marker (y-axis).

**Figure S5:** QC analysis in COVID19 datasets (Liao, M. et al. study^49^). **a**. Total counts per patient. Bar plots show the total number of UMI counts (Y-axis) per category of subjects (X-axis), shown in different color bars, from the Liao et al. study. The counts from severe and mild COVID-19 patients are depicted in orange and blue color bars while the healthy controls are green. **b**. Post-qc mitochondrial genes proportion. UMAP projection of all the post-qc filtered single cell transcriptomes from the BALF samples of all subjects in the Liao et al 2020 dataset with each cell colored by the proportion of transcripts mapped towards the human mitochondrial genes ranging from 0.0% (lowest) to 20% (highest). The color legend ranges between blue (lowest) to red (highest). **c**. Post-qc number of genes. UMAP projection of all the post-qc filtered single cell transcriptomes from the BALF samples of all subjects in the Liao et al., 2020 dataset with each cell colored by the number of detected genes ranging between 1000 to over 8000 genes. The color legend ranges between blue and red for the lowest and highest number of detected genes, respectively. **d**. Gene counts. Violin plots (with jitter points) of all the number of detected genes(Y-axis) in post-qc filtered single cell transcriptomes from the BALF samples of all subjects in the Liao, M. et al., 2020 dataset. The counts from severe and mild COVID-19 patients are depicted in orange and blue violin plots respectively while the healthy controls are green (X-axis). **e**. Mitochondrial genes fraction and cut-off. Violin plots (with jitter points) of all the proportion of mitochondrial genes (Y-axis) in pre-qc (left panel) and post-qc (right panel) filtered single cell transcriptomes from the BALF samples of all subjects in the Liao, M. et al., 2020 dataset. The proportions from severe and mild Covid19 patients are depicted in orange and blue violin plots respectively while the healthy controls are green (X-axis). **f**. Mitochondrial genes versus log total counts. Scatter plots of all the proportion of mitochondrial genes (X-axis) vs log of UMI counts in post-qc filtered single cell transcriptomes from the BALF samples of all subjects in the Liao, M. et al., 2020 dataset. Cells from severe and mild COVID-19 patients are depicted in orange and green colored points respectively while the healthy controls are blue. **g**. Log total counts versus percentage in top 100 genes. Scatter plots of all the total UMI counts in individual cell (Y-axis) versus the proportion of the genes in the top 100 most highly expressed (X-axis) in post – qc filtered single cell transcriptomes from the BALF samples of all subjects in Liao, M. et al., 2020 dataset. Cells from severe and mild COVID-19 patients are depicted in orange and blue colored points respectively while the healthy controls are green. **h**. UMI counts per single cell library. Violin plots (with jitter points) showing the total UMI counts per cell (Y-axis) in both pre – qc (left) and post-qc (right) in the three categories of subjects (X-axis) in Liao, M. et al., 2020 dataset. Cells from severe and mild COVID-19 patients are depicted in orange and blue colored violin plots respectively while the healthy controls are green. i. Percentage reads in top expressed genes per single cell library post-qc. Violin plots (with jitter points) showing the proportion of genes in the top 500 (left) and 100 (right) in post-qc filtered single cells in the three categories of subjects (X-axis) in Liao, M. et al., 2020 dataset. Cells from severe and mild COVID-19 patients are depicted in orange and green colored violin plots respectively while the healthy controls are blue.

**Figure S6:** Target genes analysis in COVID19 datasets Liao et al. study. **a**. Transcriptional state in individual subjects BALF samples. UMAP projection of individual cells transcriptional state in each subject from the Liao, M. et al., 2020 dataset. Each cell is colored in distinct color with the healthy, mild and severe covid19 subjects on the left, middle and right UMAP plots respectively. **b**. SARS-CoV-2 (nCoV) detected transcripts in individual subjects BALF samples. UMAP projection of individual cells transcriptional state in each subject from the Liao, M. et al., 2020 dataset showing the number of UMI mapped towards the SARS-CoV-2 genome in healthy, mild and severe COVID-19 subjects on the left, middle and right UMAP plots, respectively. **c**. Target genes across subjects. Violin plots (with jitter points) showing the normalized expression (Y-axis right margin) of the 17 target genes (X-axis) across the subpopulations (Y-axis left margin) in the three categories of subjects i.e. healthy(left), mild (middle) and severe (right) covid19 in Liao, M. et al., 2020 dataset.

**Figure S7:** Target genes expression in human cell atlas lung derived single cells. **a**. Target genes expression in sub-sampled 300 individual cells across Alveolar Epithelial Type1 (AET1) subpopulation (100 replications). Strip plot showing normalized expression (Y-axis) of 14 target genes (name labels above the corresponding plot) across subpopulations enriched for AET1 in which 300 individual cells were subsampled with 100 replications. The points are colored based on the lung section from which the subpopulations are derived i.e. suspected hypertensive(orange), left lobe (blue) and right lobe(green). **b**. Target genes total counts in sub-sampled 300 individual cells across AET1 subpopulation (100 replications). Strip plot showing total UMI counts (Y-axis) of 14 target genes (name labels above the corresponding plot) across subpopulations enriched for AET1 cells in which 300 individual cells were subsampled with 100 replications. The points are colored based on the lung section from which the subpopulation is derived i.e. Hypertensive(orange), left lobe (blue) and right lobe(green). **c**. Target genes proportion of cells detected in sub-sampled 300 individual cells across AET1 subpopulation (100 replications). Strip plot showing the proportion of cells(Y-axis) in which 14 target genes (name labels above the corresponding plot) were detected across subpopulations enriched for AET1 cells in which 300 individual cells were subsampled with 100 replications. The points are colored based on the lung section from which the subpopulation is derived i.e. Hypertensive (orange), left lobe (blue) and right lobe(green).

**Figure S8:** Target genes expression in COVID-19 and healthy controls at single cell resolution. **a**. Normalized mean expression in sub-sampled 300 cells across specific Ciliated Type1 subpopulation (100 replications). Strip plot showing normalized expression (Y-axis) of 22 target genes (name labels above the corresponding plot) across subpopulations enriched for Ciliated Type1 cells in which 300 individual cells were subsampled with 100 replications. The points are colored based on the subject group from which the subpopulation was derived i.e. severe covid19 (orange), mild covid19 (blue) and healthy controls (green).

**Figure S9:** Cell types enriched in subpopulations of BALF samples from healthy (a), mild (b) and severe COVID-29 cases.

## Methods

### Identification of short linear motifs (SLIMs) in SARS-CoV-1 and 2

The Eukaryotic Linear Motif (ELM) resource server ^23,24^ was used to scan the SARS-CoV-1 and 2 spike protein sequence for candidate and known functional motifs. To minimize false positive short linear motifs (SLIMs), we used filters that include contextual information on the location of the motif such that motifs occurring within globular domains are excluded. Detected ITIM sequences were manually verified to be consistent with ITIM regular expression (ILV)-x-x-Y-x-(LV). ITIM containing proteins in the human proteome were obtained from a previous study^30^.

### Selection of candidate genes

Candidate genes for transcriptome analysis were selected based on their roles in i) SARS-CoV-2 cell entry pathways (*ACE2, TMPRSS2, NRP1, NRP2*)^50–54^; ii) ADE via Fc receptor binding and/or containing ITIM/ITAM sequences (*FCGR2A*^17,18^, *FCGR2B*^7^, *LILRB1*^19,20^, *NKG2A*)^55^; iii) disruption of ITIM-dependent signaling by synthetic ITIM containing peptides (*MyD88* and *CD300LF/IREM-1*)^21,22^; iv) F_c_ core fucosylation which enhances ADE (*Fut8*)^56,57^; iv) innate immunity (*IFNG* and *IFNB*); v) functional similarity to the spike protein (syncytin/*ERVW1* involved in formation of the placenta through cell-to-cell fusion under the control of LIF^101,102^). Like the S protein, syncytin is a class I fusion protein^103^ and its ligand in mice (Ly6E) was recently shown to impair SARS-CoV-2 membrane fusion^104^. Furthermore, HERV expression impacts innate immunity to exogenous viruses and some ERVs have been co-opted as regulators of innate immunity in non-human primates^83,105^.

### Datasets

We used published single cell RNA-seq (scRNA-seq) data, for different tissues/organs, from the human cell atlas data portal (https://data.humancellatlas.org/)^47^. For the normal lung expression data we used datasets from the Madissoon et al study^48^ while the bronchial alveolar lavage fluid (BALF) derived cells from COVID-19 patients from the Liao et al., 2020 study^49^ (GEO accession number GSE145926) used in the analysis. In both studies we only used cells with at least 300 genes detected, mitochondrial gene fraction<20% and at least 10,000 mapped UMI per cell for the quality control, target genes mean expression and initial exploratory analysis.

### Cell subpopulations

We used single cells that had at least 1000 genes detected, mitochondrial gene fraction<20% and at least 10,000 mapped UMI per cell. Each cell’s total counts were normalized to 10,000, then log transformed, and significantly highly variable genes identified using in Seurat’s mean expression weighted dispersion Z -scores as implemented in Scanpy module (version 1.5.1) (Wolf, F.A., Angerer, P. & Theis, F.J., 2018). Dimensionality reduction was carried out using PCA and the top 20 components used as input to Louvain algorithm clustering, with the resolution set to 1.0 and other default parameters. The neighborhood graphs were based on UMAP embedding and Euclidean distance.

### Cell type detection

Here we compiled a set of markers for each organ/tissue cell type as described in the UCSC cell browser (https://cells.ucsc.edu/). For each subpopulation calculated the overlap score between its top 100 markers and each cell type markers using Scanpy’s marker_gene_overlap algorithm^62^ with normalization set to the ‘reference’ set of markers.

### Differential gene expression analysis

Since the number of cells varied for each subpopulation, we subsampled 300 cells from each subpopulation and performed differential gene expression using Welch – t test using the genes counts with negative binomial noise model. 100 replications were carried out for each pairwise comparison between subpopulations.

### Code and datasets

All the results datasets and the accompanying code can be accessed as jupyter notebook from GitHub at https://github.com/SiwoResearch/SARS-CoV-2-Immunopathology

## Conflicts of interest declaration

The authors declare no conflict of interest.

## Acknowledgements

We are grateful to Dr. Carol Sibley (University of Washington) for the insightful discussions and feedback on this work.

## Notes

https://github.com/SiwoResearch/SARS-CoV-2-Immunopathology

### Competing Interest Statement

The authors have declared no competing interest.

## References

1. Hu, B., Huang, S. & Yin, L. The cytokine storm and COVID-19. Journal of Medical Virology (2021) doi:10.1002/jmv.26232.

2. Sarma, A. et al. COVID-19 ARDS is characterized by a dysregulated host response that differs from cytokine storm and is modified by dexamethasone. Res. Sq. (2021) doi:10.21203/rs.3.rs-141578/v1.

3. Bastard, P. et al. Autoantibodies against type I IFNs in patients with life-threatening COVID-19. Science (80-.). (2020) doi:10.1126/science.abd4585.

4. Woodruff, M. C., Ramonell, R. P., Lee, F. E. H. & Sanz, I. Clinically identifiable autoreactivity is common in severe SARS-CoV-2 infection. medRxiv (2020) doi:10.1101/2020.10.21.20216192.

5. Wang, E. Y. et al. Diverse functional autoantibodies in patients with COVID-19. medRxiv (2020) doi:10.1101/2020.12.10.20247205.

6. Kuroki, K., Furukawa, A. & Maenaka, K. Molecular recognition of paired receptors in the immune system. Frontiers in Microbiology (2012) doi:10.3389/fmicb.2012.00429.

7. Kane, B. A., Bryant, K. J., McNeil, H. P. & Tedla, N. T. Termination of immune activation: An essential component of healthy host immune responses. Journal of Innate Immunity (2014) doi:10.1159/000363449.

8. Ivashkiv, L. B. How ITAMs inhibit signaling. Science Signaling (2011) doi:10.1126/scisignal.2001917.

9. Ong, E. Z., Chan, K. R. & Ooi, E. E. Viral Manipulation of Host Inhibitory Receptor Signaling for Immune Evasion. PLoS Pathogens (2016) doi:10.1371/journal.ppat.1005776.

10. Van Avondt, K., van Sorge, N. M. & Meyaard, L. Bacterial Immune Evasion through Manipulation of Host Inhibitory Immune Signaling. PLoS Pathogens (2015) doi:10.1371/journal.ppat.1004644.

11. Davey, N. E., Travé, G. & Gibson, T. J. How viruses hijack cell regulation. Trends in Biochemical Sciences (2011) doi:10.1016/j.tibs.2010.10.002.

12. Via, A., Uyar, B., Brun, C. & Zanzoni, A. How pathogens use linear motifs to perturb host cell networks. Trends in Biochemical Sciences (2015) doi:10.1016/j.tibs.2014.11.001.

13. Yang, C. W. & Shi, Z. L. Uncovering potential host proteins and pathways that may interact with eukaryotic short linear motifs in viral proteins of MERS, SARS and SARS2 coronaviruses that infect humans. PLoS One (2021) doi:10.1371/journal.pone.0246150.

14. Chemes, L. B., de Prat-Gay, G. & Sánchez, I. E. Convergent evolution and mimicry of protein linear motifs in host-pathogen interactions. Current Opinion in Structural Biology (2015) doi:10.1016/j.sbi.2015.03.004.

15. Wang, Q. et al. Immunodominant SARS coronavirus epitopes in humans elicited both enhancing and neutralizing effects on infection in non-human primates. ACS Infect. Dis. (2016) doi:10.1021/acsinfecdis.6b00006.

16. Liu, L. et al. Anti-spike IgG causes severe acute lung injury by skewing macrophage responses during acute SARS-CoV infection. JCI insight (2019) doi:10.1172/jci.insight.123158.

17. Marovich, M. A., Boonnak, K., Slike, B. M. & Donofrio, G. C. Dengue Virus Infection Differentially Influence Antibody-Mediated RII Cytoplasmic Domains γ Human Fc Human FcgRII Cytoplasmic Domains Differentially Influence Antibody-Mediated Dengue Virus Infection. J. Immunol. (2013) doi:10.4049/jimmunol.1203052.

18. Moi, M. L., Lim, C. K., Takasaki, T. & Kurane, I. Involvement of the Fcγ receptor IIA cytoplasmic domain in antibody-dependent enhancement of dengue virus infection. J. Gen. Virol. (2010) doi:10.1099/vir.0.014829-0.

19. Ong, E. Z. et al. Dengue virus compartmentalization during antibody-enhanced infection. Sci. Rep. (2017) doi:10.1038/srep40923.

20. Chan, K. R. et al. Leukocyte immunoglobulin-like receptor B1 is critical for antibody-dependent dengue. Proc. Natl. Acad. Sci. U. S. A. (2014) doi:10.1073/pnas.1317454111.

21. Lee, S. M., Suk, K. & Lee, W. H. Synthetic peptides containing ITIM-like sequences of IREM-1 (CD300F) differentially regulate MyD88 and TRIF-mediated TLR signalling through activation of SHP and/or PI3K. Clin. Exp. Immunol. (2012) doi:10.1111/j.1365-2249.2011.04528.x.

22. Lee, S.-M., Kim, E.-J., Suk, K. & Lee, W.-H. CD300F Blocks Both MyD88 and TRIF-Mediated TLR Signaling through Activation of Src Homology Region 2 Domain-Containing Phosphatase 1. J. Immunol. (2011) doi:10.4049/jimmunol.1002184.

23. Dinkel, H. et al. ELM -The database of eukaryotic linear motifs. Nucleic Acids Res. (2012) doi:10.1093/nar/gkr1064.

24. Kumar, M. et al. ELM-the eukaryotic linear motif resource in 2020. Nucleic Acids Res. (2020) doi:10.1093/nar/gkz1030.

25. Mészáros, B. et al. Short linear motif candidates in the cell entry system used by SARS-CoV-2 and their potential therapeutic implications. Sci. Signal. (2021) doi:10.1126/SCISIGNAL.ABD0334.

26. Korber, B. et al. Tracking Changes in SARS-CoV-2 Spike: Evidence that D614G Increases Infectivity of the COVID-19 Virus. Cell (2020) doi:10.1016/j.cell.2020.06.043.

27. Korber, B. et al. Spike mutation pipeline reveals the emergence of a more transmissible form of SARS-CoV-2. bioRxiv (2020) doi:10.1101/2020.04.29.069054.

28. Grubaugh, N. D., Hanage, W. P. & Rasmussen, A. L. Making Sense of Mutation: What D614G Means for the COVID-19 Pandemic Remains Unclear. Cell (2020) doi:10.1016/j.cell.2020.06.040.

29. Oliver, S. L. et al. An immunoreceptor tyrosine-based inhibition motif in varicella-zoster virus glycoprotein B regulates cell fusion and skin pathogenesis. Proc. Natl. Acad. Sci. U. S. A. (2013) doi:10.1073/pnas.1216985110.

30. Staub, E., Rosenthal, A. & Hinzmann, B. Systematic identification of immunoreceptor tyrosine-based inhibitory motifs in the human proteome. Cell. Signal. (2004) doi:10.1016/j.cellsig.2003.08.013.

31. Quinton, L. J. et al. Leukemia Inhibitory Factor Signaling Is Required for Lung Protection during Pneumonia. J. Immunol. (2012) doi:10.4049/jimmunol.1200256.

32. Foronjy, R. F., Dabo, A. J., Cummins, N. & Geraghty, P. Leukemia inhibitory factor protects the lung during respiratory syncytial viral infection. BMC Immunol. (2014) doi:10.1186/s12865-014-0041-4.

33. Poon, J. et al. Cigarette Smoke Exposure Reduces Leukemia Inhibitory Factor Levels During Respiratory Syncytial Viral Infection. in (2019). doi:10.1164/ajrccm-conference.2019.199.1_meetingabstracts.a4517.

34. Traber, K. E. et al. Myeloid-epithelial cross talk coordinates synthesis of the tissue-protective cytokine leukemia inhibitory factor during pneumonia. Am. J. Physiol. -Lung Cell. Mol. Physiol. (2017) doi:10.1152/ajplung.00482.2016.

35. Jorens, P. G. et al. High levels of leukaemia inhibitory factor in ARDS. Cytokine (1996) doi:10.1006/cyto.1996.9999.

36. Gruson, D. et al. Sequential production of leukaemia inhibitory factor by blood cell culture in patients with ARDS. Intensive Care Med. (1998) doi:10.1007/s001340050582.

37. Wang, D. et al. Clinical Characteristics of 138 Hospitalized Patients With 2019 Novel Coronavirus–Infected Pneumonia in Wuhan, China. JAMA (2020) doi:10.1001/jama.2020.1585.

38. Fan, E. et al. COVID-19-associated acute respiratory distress syndrome: is a different approach to management warranted? The Lancet Respiratory Medicine (2020) doi:10.1016/S2213-2600(20)30304-0.

39. Daniloski, Z., Guo, X. & Sanjana, N. E. The D614G mutation in SARS-CoV-2 Spike increases transduction of multiple human cell types. bioRxiv (2020) doi:10.1101/2020.06.14.151357.

40. Zhang, L. et al. The D614G mutation in the SARS-CoV-2 spike protein reduces S1 shedding and increases infectivity. bioRxiv (2020) doi:10.1101/2020.06.12.148726.

41. Hu, J. et al. The D614G mutation of SARS-CoV-2 spike protein enhances viral infectivity and decreases neutralization sensitivity to individual convalescent sera. bioRxiv (2020).

42. Mansbach, R. A. et al. The SARS-CoV-2 Spike Variant D614G Favors an Open Conformational State. Biophys. J. (2021) doi:10.1016/j.bpj.2020.11.1904.

43. Daniloski, Z. et al. The spike d614g mutation increases sars-cov-2 infection of multiple human cell types. Elife (2021) doi:10.7554/eLife.65365.

44. Hou, Y. J. et al. SARS-CoV-2 D614G variant exhibits efficient replication ex vivo and transmission in vivo. Science (80-.). (2021) doi:10.1126/science.abe8499.

45. Zhang, J. et al. Structural impact on SARS-CoV-2 spike protein by D614G substitution. Science (80-.). (2021) doi:10.1126/science.abf2303.

46. Plante, J. A. et al. Spike mutation D614G alters SARS-CoV-2 fitness. Nature (2020) doi:10.1038/s41586-020-2895-3.

47. Regev, A. et al. The human cell atlas. Elife (2017) doi:10.7554/eLife.27041.

48. Madissoon, E. et al. ScRNA-seq assessment of the human lung, spleen, and esophagus tissue stability after cold preservation. Genome Biol. (2019) doi:10.1186/s13059-019-1906-x.

49. Liao, M. et al. Single-cell landscape of bronchoalveolar immune cells in patients with COVID-19. Nat. Med. (2020) doi:10.1038/s41591-020-0901-9.

50. George Sakoulas, M. ACE2 Is the SARS-CoV-2 Receptor Required for Cell Entry. NEJM J. Watch (2020).

51. Scialo, F. et al. ACE2: The Major Cell Entry Receptor for SARS-CoV-2. Lung (2020) doi:10.1007/s00408-020-00408-4.

52. Hoffmann, M. et al. SARS-CoV-2 Cell Entry Depends on ACE2 and TMPRSS2 and Is Blocked by a Clinically Proven Protease Inhibitor. Cell (2020) doi:10.1016/j.cell.2020.02.052.

53. Cantuti-Castelvetri, L. et al. Neuropilin-1 facilitates SARS-CoV-2 cell entry and infectivity. Science (80-.). (2020) doi:10.1126/science.abd2985.

54. Daly, J. L. et al. Neuropilin-1 is a host factor for SARS-CoV-2 infection. Science (80-.). (2020) doi:10.1126/science.abd3072.

55. Kabat, J., Borrego, F., Brooks, A. & Coligan, J. E. Role That Each NKG2A Immunoreceptor Tyrosine-Based Inhibitory Motif Plays in Mediating the Human CD94/NKG2A Inhibitory Signal. J. Immunol. (2002) doi:10.4049/jimmunol.169.4.1948.

56. Larsen, M. D. et al. Afucosylated IgG characterizes enveloped viral responses and correlates with COVID-19 severity. Science (80-.). (2021) doi:10.1126/science.abc8378.

57. Wang, T. T. et al. IgG antibodies to dengue enhanced for FcγRIIIA binding determine disease severity. Science (2017) doi:10.1126/science.aai8128.

58. Sungnak, W. et al. SARS-CoV-2 entry factors are highly expressed in nasal epithelial cells together with innate immune genes. Nat. Med. (2020) doi:10.1038/s41591-020-0868-6.

59. Singh, M., Bansal, V. & Feschotte, C. A Single-Cell RNA Expression Map of Human Coronavirus Entry Factors. Cell Rep. (2020) doi:10.1016/j.celrep.2020.108175.

60. McInnes, L., Healy, J. & Melville, J. UMAP: Uniform manifold approximation and projection for dimension reduction. arXiv (2018).

61. Blondel, V. D., Guillaume, J. L., Lambiotte, R. & Lefebvre, E. Fast unfolding of communities in large networks. J. Stat. Mech. Theory Exp. (2008) doi:10.1088/1742-5468/2008/10/P10008.

62. Wolf, F. A., Angerer, P. & Theis, F. J. SCANPY: Large-scale single-cell gene expression data analysis. Genome Biol. (2018) doi:10.1186/s13059-017-1382-0.

63. Cañadas, I. et al. Tumor innate immunity primed by specific interferon-stimulated endogenous retroviruses. Nat. Med. (2018) doi:10.1038/s41591-018-0116-5.

64. Chiappinelli, K. B. et al. Inhibiting DNA Methylation Causes an Interferon Response in Cancer via dsRNA Including Endogenous Retroviruses. Cell (2017) doi:10.1016/j.cell.2017.03.036.

65. Dhanwani, R. et al. Cellular sensing of extracellular purine nucleosides triggers an innate IFN-β response. Sci. Adv. (2020) doi:10.1126/sciadv.aba3688.

66. Menezes, S. M., Braz, M., Llorens-Rico, V., Wauters, J. & Van Weyenbergh, J. Endogenous IFNβ expression predicts outcome in critical patients with COVID-19. The Lancet. Microbe (2021) doi:10.1016/S2666-5247(21)00063-X.

67. Maucourant, C. et al. Natural killer cell immunotypes related to COVID-19 disease severity. Sci. Immunol. (2020) doi:10.1126/SCIIMMUNOL.ABD6832.

68. Mathew, D. et al. Deep immune profiling of COVID-19 patients reveals distinct immunotypes with therapeutic implications. Science (80-.). (2020) doi:10.1126/SCIENCE.ABC8511.

69. Hirsch, I., Janovec, V., Stranska, R. & Bendriss-Vermare, N. Cross talk between inhibitory immunoreceptor tyrosine-based activation motif-signaling and toll-like receptor pathways in macrophages and dendritic cells. Frontiers in Immunology (2017) doi:10.3389/fimmu.2017.00394.

70. Poe, J. C. & Tedder, T. F. CD22 and Siglec-G in B cell function and tolerance. Trends in Immunology (2012) doi:10.1016/j.it.2012.04.010.

71. Metcalfe, S. M. LIF and the lung’s stem cell niche: Is failure to use LIF to protect against COVID-19 a grave omission in managing the pandemic? Future Virology (2020) doi:10.2217/fvl-2020-0340.

72. Rose-John, S. nterleukin-6 family cytokines. Cold Spring Harb. Perspect. Biol. (2018) doi:10.1101/cshperspect.a028415.

73. Giese, B. et al. Dimerization of the cytokine receptors gp130 and LIFR analysed in single cells. J. Cell Sci. (2005) doi:10.1242/jcs.02628.

74. Hiemstra, P. S., McCray, P. B. & Bals, R. The innate immune function of airway epithelial cells in inflammatory lung disease. Eur. Respir. J. (2015) doi:10.1183/09031936.00141514.

75. European Medicines Agency. EMA endorses use of dexamethasone in COVID-19 patients on oxygen or mechanical ventilation. 18/9 (2020).

76. Stauffer, W. M., Alpern, J. D. & Walker, P. F. COVID-19 and Dexamethasone. JAMA (2020) doi:10.1001/jama.2020.13170.

77. Liu, X. G. et al. High-dose dexamethasone shifts the balance of stimulatory and inhibitory Fcγ receptors on monocytes in patients with primary immune thrombocytopenia. Blood (2011) doi:10.1182/blood-2010-07-295477.

78. Silwal, P. et al. Dexamethasone induces FCRIIB expression in RBL-2H3 cells. Korean J. Physiol. Pharmacol. (2012) doi:10.4196/kjpp.2012.16.6.393.

79. Stanciu, L. A. et al. Expression of programmed death-1 ligand (PD-L) 1, PD-L2, B7-H3, and inducible costimulator ligand on human respiratory tract epithelial cells and regulation by respiratory syncytial virus and type 1 and 2 cytokines. J. Infect. Dis. (2006) doi:10.1086/499275.

80. Thai, L. M., Ashman, L. K., Harbour, S. N., Hogarth, P. M. & Jackson, D. E. Physical proximity and functional interplay of PECAM-1 with the Fc receptor FcγRIIa on the platelet plasma membrane. Blood (2003) doi:10.1182/blood-2003-02-0496.

81. Schultz, N. H. et al. Thrombosis and Thrombocytopenia after ChAdOx1 nCoV-19 Vaccination. N. Engl. J. Med. 0, null.

82. Greinacher, A. et al. Thrombotic Thrombocytopenia after ChAdOx1 nCov-19 Vaccination. N. Engl. J. Med. 0, null.

83. Chuong, E. B., Elde, N. C. & Feschotte, C. Regulatory evolution of innate immunity through co-option of endogenous retroviruses. Science (80-.). (2016) doi:10.1126/science.aad5497.

84. Zhang, L. et al. SARS-CoV-2 RNA reverse-transcribed and integrated into the human genome. bioRxiv (2020) doi:10.1101/2020.12.12.422516.

85. Pavlicek, A., Paces, J., Elleder, D. & Hejnar, J. Processed Pseudogenes of Human Endogenous Retroviruses Generated by LINEs: Their Integration, Stability, and Distribution. Genome Res. (2002) doi:10.1101/gr.216902.

86. Dupressoir, A., Lavialle, C. & Heidmann, T. From ancestral infectious retroviruses to bona fide cellular genes: Role of the captured syncytins in placentation. Placenta (2012) doi:10.1016/j.placenta.2012.05.005.

87. Kremer, D. et al. PHERV-W envelope protein fuels microglial cell-dependent damage of myelinated axons in multiple sclerosis. Proc. Natl. Acad. Sci. U. S. A. (2019) doi:10.1073/pnas.1901283116.

88. Curtin, F. et al. A new therapeutic approach for type 1 diabetes: Rationale for GNbAC1, an anti-HERV-W-Env monoclonal antibody. Diabetes, Obesity and Metabolism (2018) doi:10.1111/dom.13357.

89. Niegowska, M. et al. Anti-HERV-W Env antibodies are correlated with seroreactivity against Mycobacterium avium subsp. paratuberculosis in children and youths at T1D risk. Sci. Rep. (2019) doi:10.1038/s41598-019-42788-5.

90. Rosas, I. O. et al. Tocilizumab in Hospitalized Patients with Severe Covid-19 Pneumonia. N. Engl. J. Med. (2021) doi:10.1056/nejmoa2028700.

91. Interleukin-6 Receptor Antagonists in Critically Ill Patients with Covid-19. N. Engl. J. Med. (2021) doi:10.1056/nejmoa2100433.

92. Fioravante, M. et al. Inhibition of hypothalamic leukemia inhibitory factor exacerbates diet-induced obesity phenotype. J. Neuroinflammation (2017) doi:10.1186/s12974-017-0956-9.

93. Wang, T. et al. Effects of leukemia inhibitory factor receptor on the adipogenic differentiation of human bone marrow mesenchymal stem cells. Mol. Med. Rep. (2019) doi:10.3892/mmr.2019.10140.

94. Wang, L., Wu, Q., Wang, R. Q., Wang, R. Z. & Wang, J. Protection of leukemia inhibitory factor against high-glucose-induced human retinal endothelial cell dysfunction. Arch. Physiol. Biochem. (2020) doi:10.1080/13813455.2020.1792506.

95. López, N., Varo, N., Díez, J. & Fortuño, M. A. Loss of myocardial LIF receptor in experimental heart failure reduces cardiotrophin-1 cytoprotection. A role for neurohumoral agonists? Cardiovasc. Res. (2007) doi:10.1016/j.cardiores.2007.04.025.

96. Chen, K., Zhou, M., Wang, X., Li, S. & Yang, D. The Role of Myokines and Adipokines in Hypertension and Hypertension-related Complications. Hypertens. Res. (2019) doi:10.1038/s41440-019-0266-y.

97. Davis, S. M. et al. Leukemia Inhibitory Factor Protects Neurons from Ischemic Damage via Upregulation of Superoxide Dismutase 3. Mol. Neurobiol. (2017) doi:10.1007/s12035-015-9587-2.

98. Yoshino, J., Monkawa, T., Tsuji, M., Hayashi, M. & Saruta, T. Leukemia Inhibitory Factor Is Involved in Tubular Regeneration after Experimental Acute Renal Failure. J. Am. Soc. Nephrol. (2003) doi:10.1097/01.ASN.0000101180.96787.02.

99. Lopez-Arenas, E., Mackay-Sim, A., Bacigalupo, J. & Sulz, L. Leukaemia Inhibitory Factor Stimulates Proliferation of Olfactory Neuronal Progenitors via Inducible Nitric Oxide Synthase. PLoS One (2012) doi:10.1371/journal.pone.0045018.

100. Nan, B., Getchell, M. L., Partin, J. V. & Getchell, T. V. Leukemia inhibitory factor, interleukin-6, and their receptors are expressed transiently in the olfactory mucosa after target ablation. J. Comp. Neurol. (2001) doi:10.1002/cne.1193.

101. Kojima, K. et al. Expression of leukaemia inhibitory factor (LIF) receptor in human placenta: A possible role for LIF in the growth and differentiation of trophoblasts. Mol. Hum. Reprod. (1995) doi:10.1093/molehr/1.5.249.

102. Suman, P., Malhotra, S. S. & Gupta, S. K. LIF-STAT signaling and trophoblast biology. JAK-STAT (2013) doi:10.4161/jkst.25155.

103. Vance, T. D. R. & Lee, J. E. Virus and eukaryote fusogen superfamilies. Curr. Biol. (2020) doi:10.1016/j.cub.2020.05.029.

104. Pfaender, S. et al. LY6E impairs coronavirus fusion and confers immune control of viral disease. Nat. Microbiol. (2020) doi:10.1038/s41564-020-0769-y.

105. Katzourakis, A. & Aswad, A. Evolution: Endogenous viruses provide shortcuts in antiviral immunity. Current Biology (2016) doi:10.1016/j.cub.2016.03.072.

