## Supplementary Figure 1 for "Molecular relationships between SARS-CoV-2 Spike protein and LIFR, a pneumonia protective IL-6 family cytokine receptor"

### Figure S1 : Immunopathology analysis in human cell atlas datasets

A. Track plot of target genes across cells across normal tissues in HCA at single cell resolution

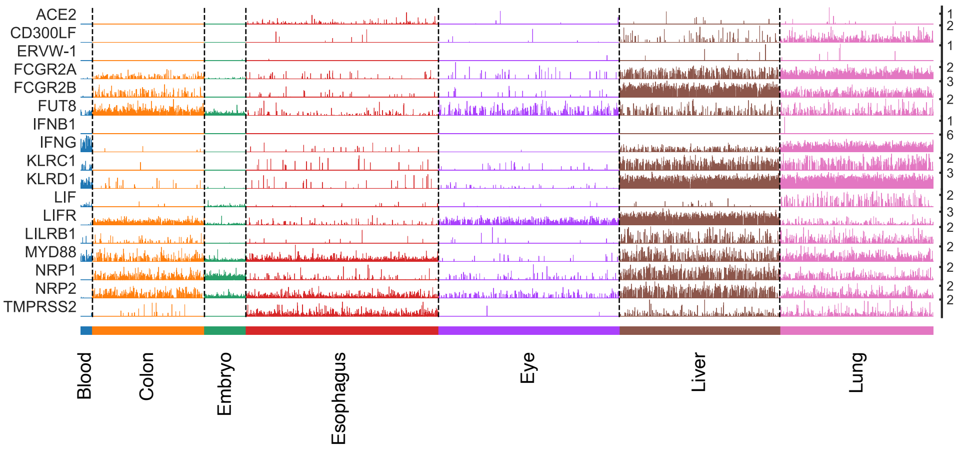


B. Target genes detection rate in single cells normal across tissues

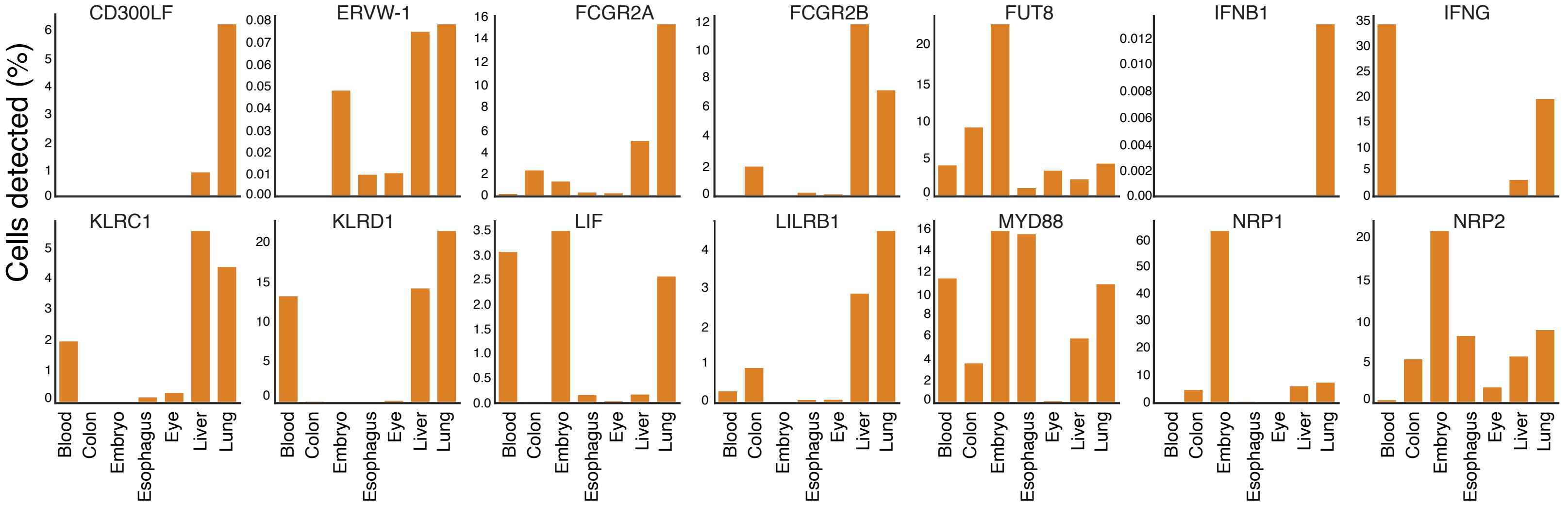


C. Target genes expression in lung-derived cells (Madissoon et al. dataset)

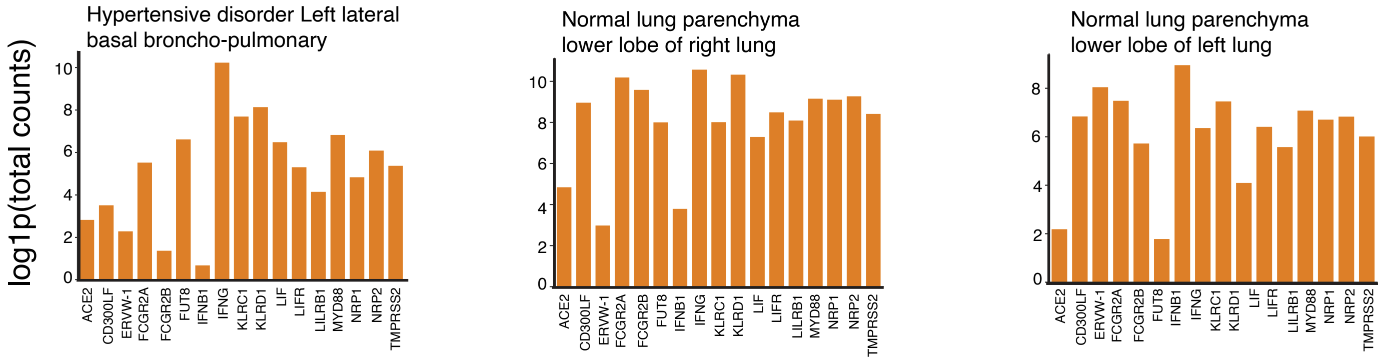


D. Target genes detection rate in lung-derived cells (Madissoon et al. dataset)

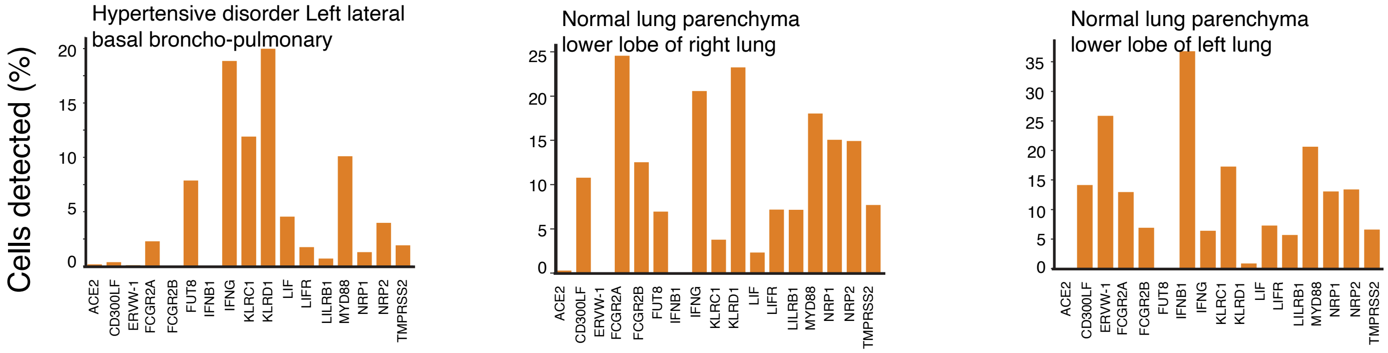
