## Supplementary figures and images for "Molecular relationships between SARS-CoV-2 Spike protein and LIFR, a pneumonia protective IL-6 family cytokine receptor"

### Supplementary Figure 2

**Figure S2: Expression of immunopathology genes across human tissues in HCA data**

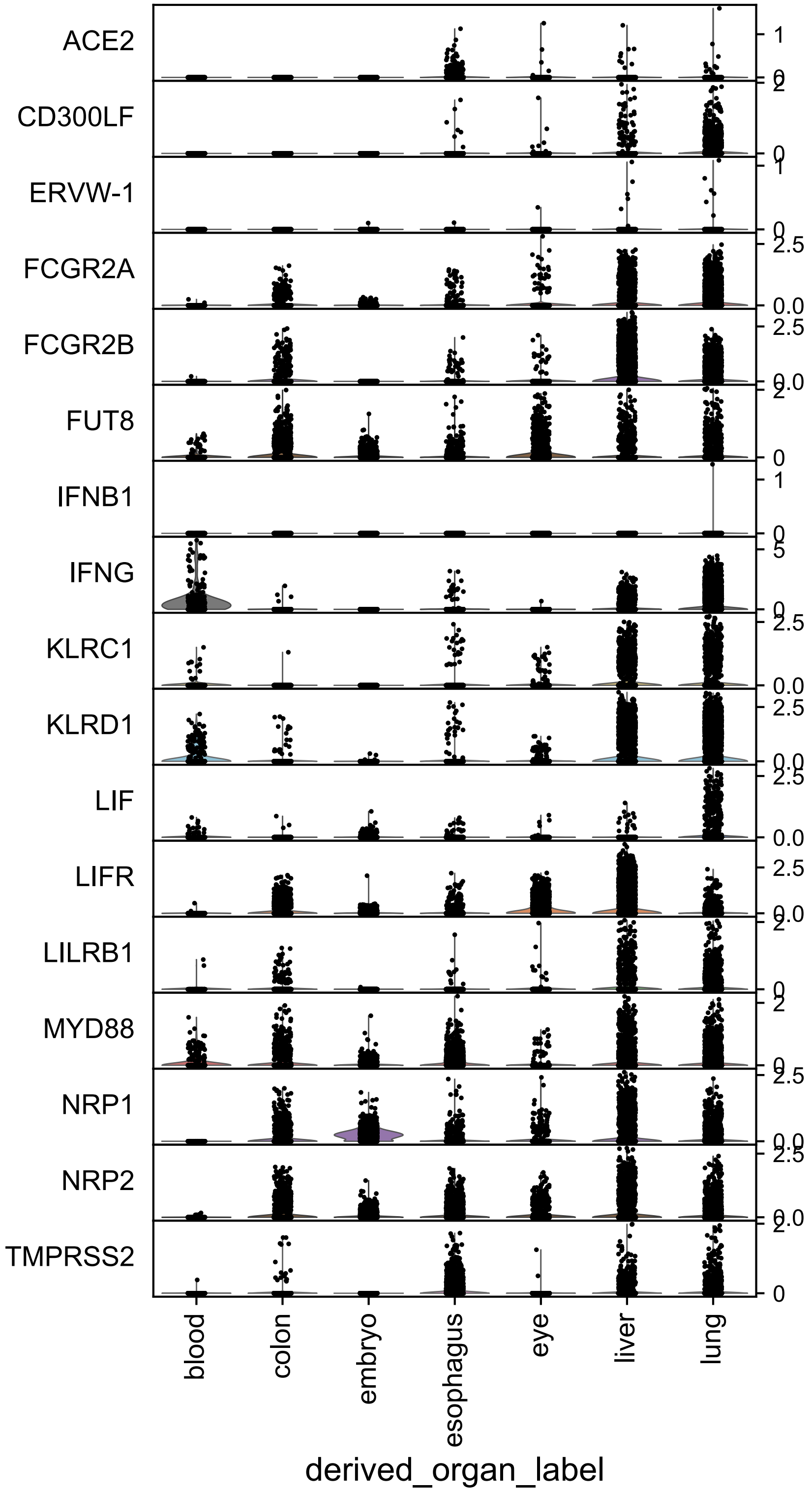

### Supplementary Figure 5

#### Figure S5 : QC analysis in COVID19 datasets (Liao, M. et al. dataset)

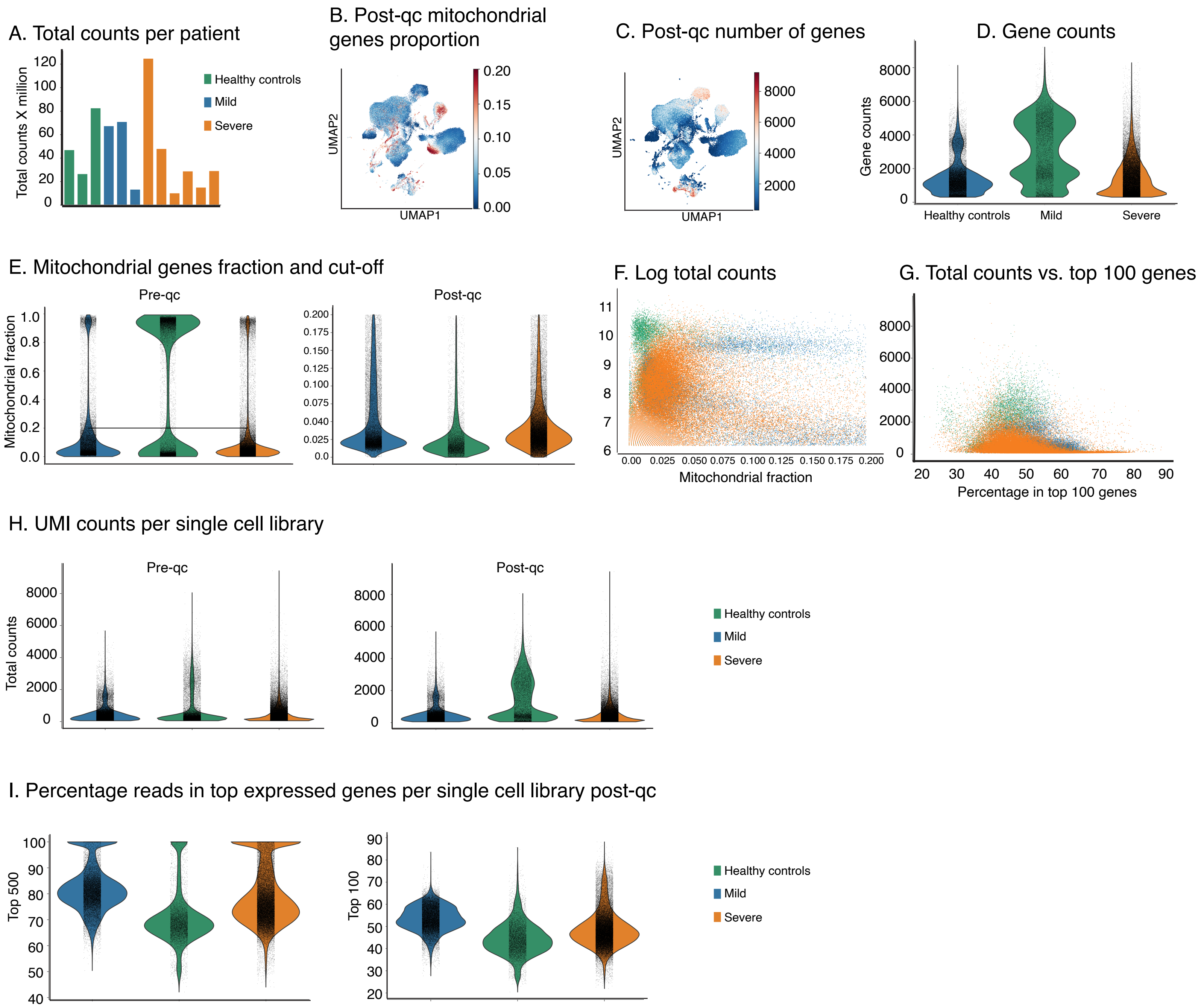
