## Supplementary Figure 3 for "Molecular relationships between SARS-CoV-2 Spike protein and LIFR, a pneumonia protective IL-6 family cytokine receptor"

### Figure S3: Target genes expression within HCA lung (Madissoon et al. dataset)

#### Target genes in individual lung cells

##### A. Hypertensive disoder subpopulations

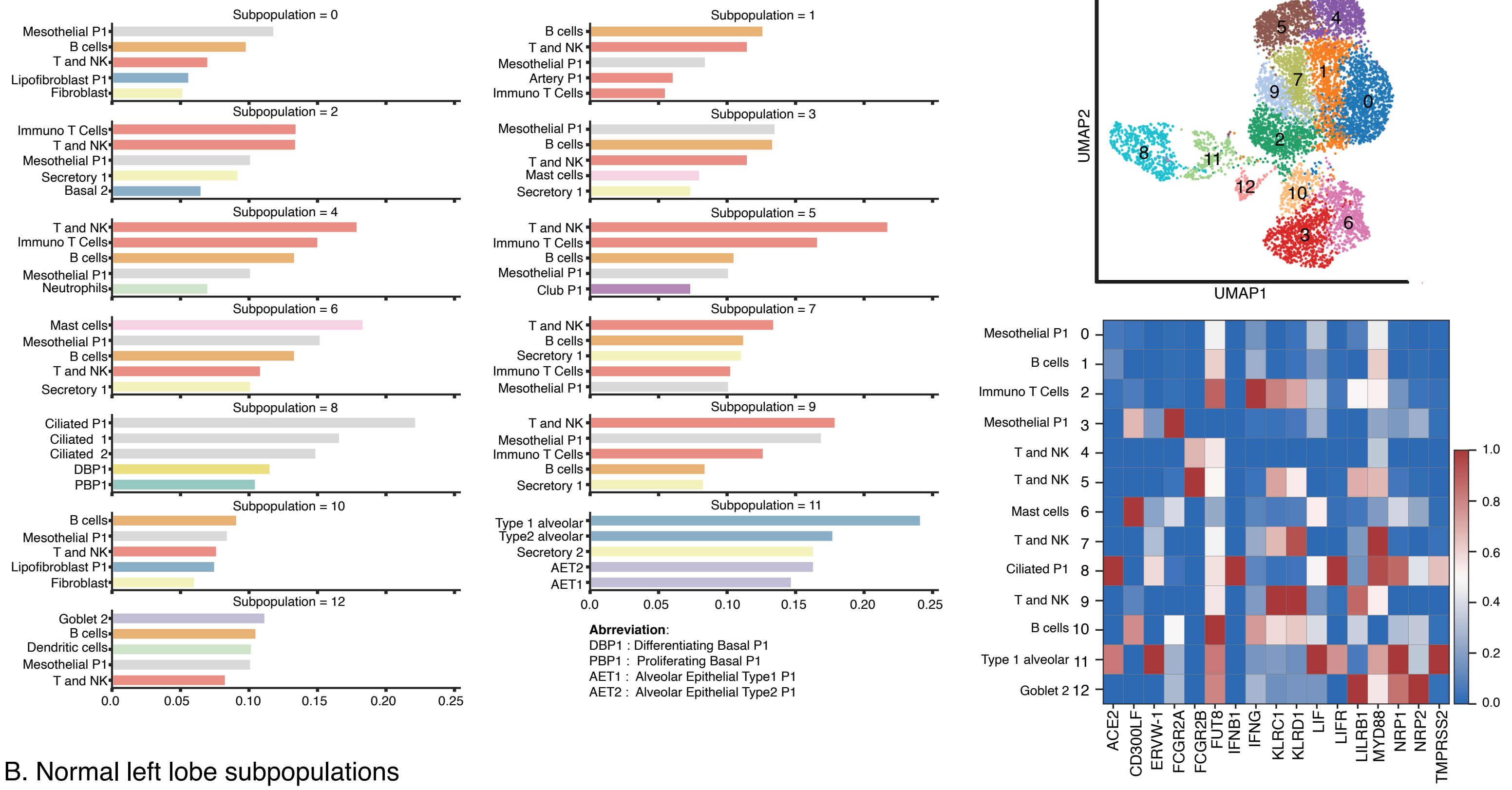


##### B. Normal left lobe subpopulations

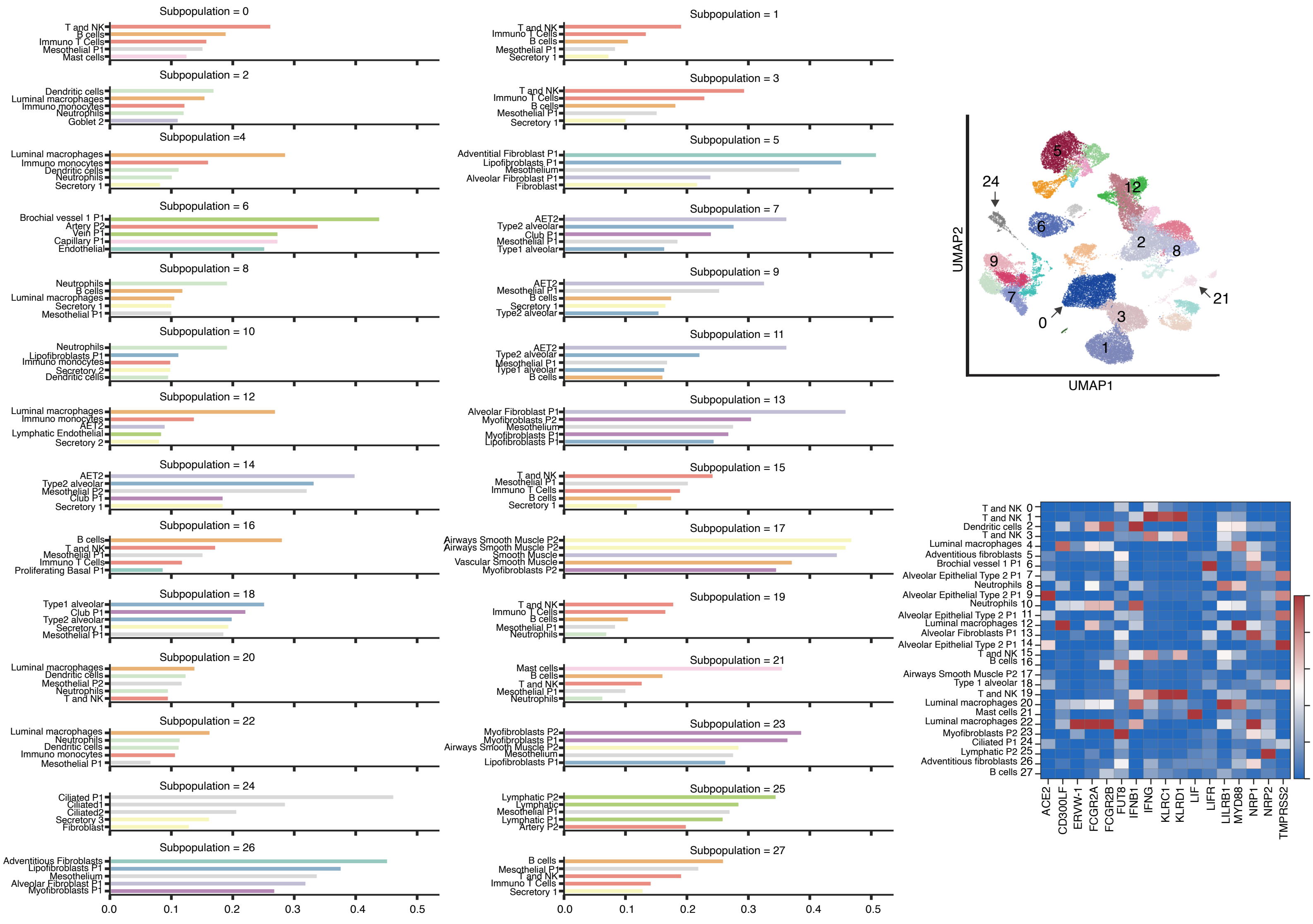
