## Supplementary Figure 4 for "Molecular relationships between SARS-CoV-2 Spike protein and LIFR, a pneumonia protective IL-6 family cytokine receptor"

### Figure S4 : Target genes expression within HCA lung(Madissoon et al. dataset)

Normal right lobe subpopulations

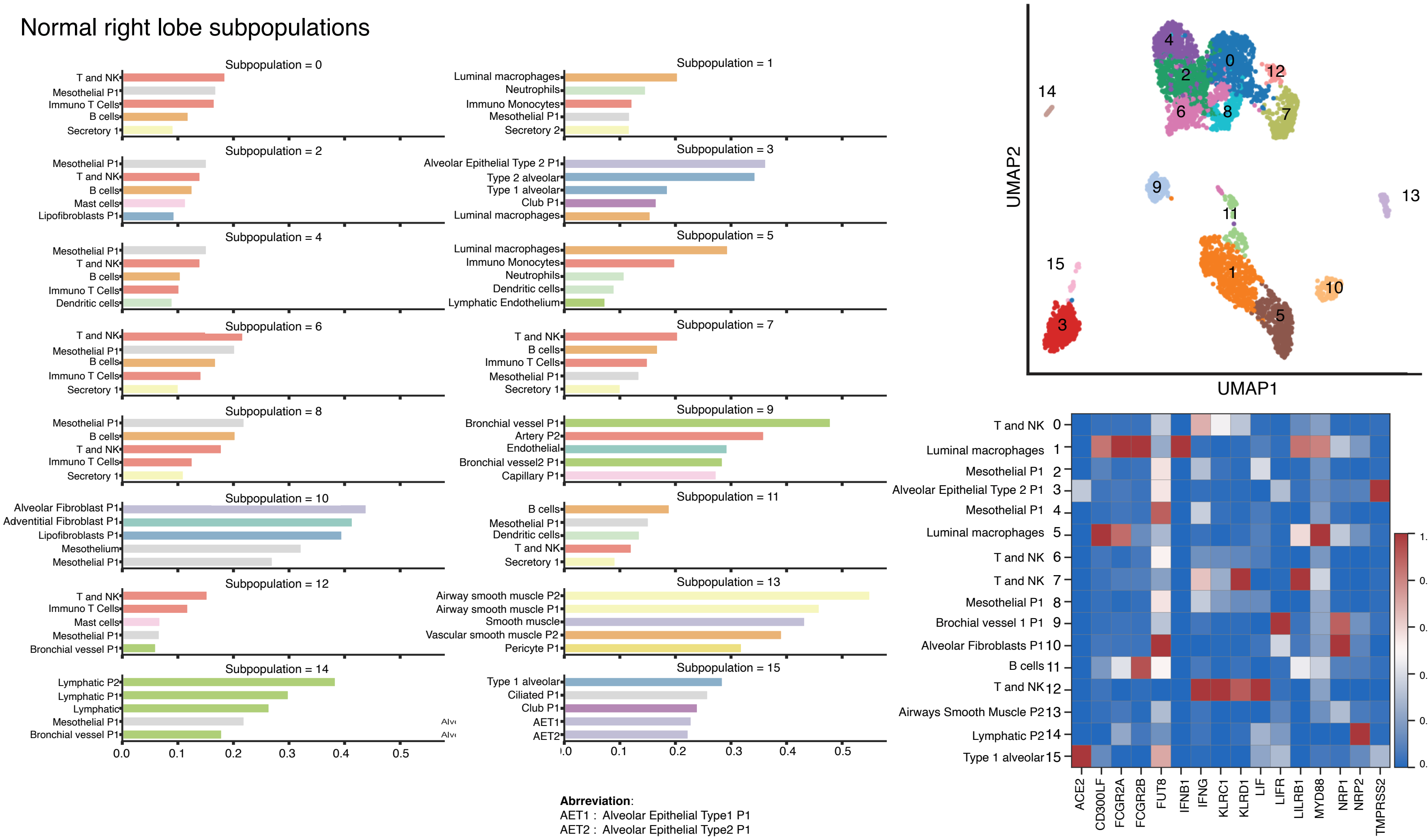


**Abbreviation**:
AET1 : Alveolar Epithelial Type1 P1
AET2 : Alveolar Epithelial Type2 P1
