## Supplementary Figure 6 for "Molecular relationships between SARS-CoV-2 Spike protein and LIFR, a pneumonia protective IL-6 family cytokine receptor"

### Figure S6 : Target genes analysis in COVID19 datasets (Liao, M. et al. dataset)

A. Transcriptional state in individual subjects BALF samples

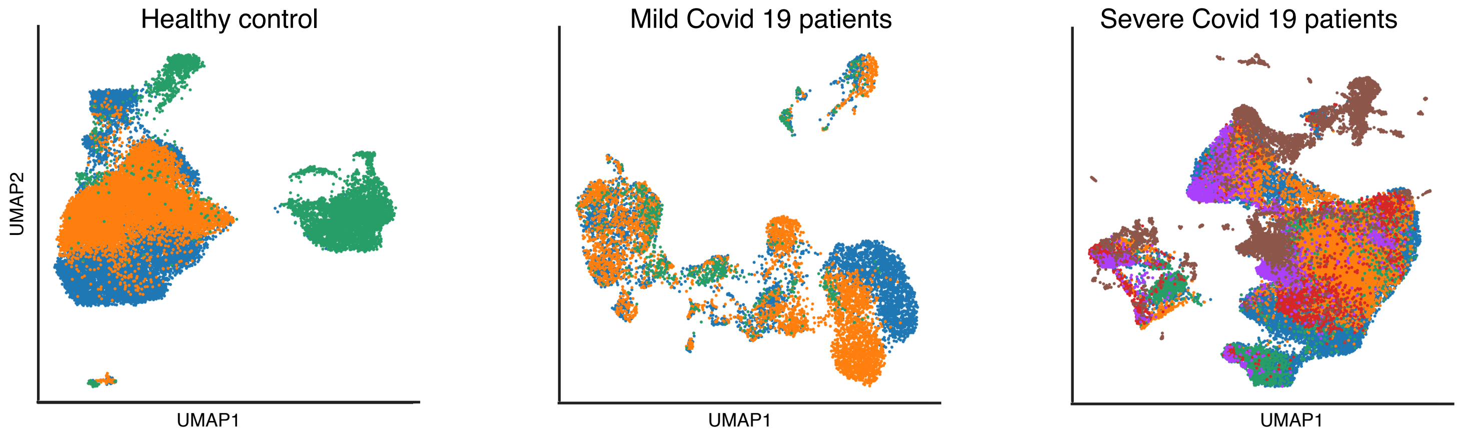


B. nCoV detected transcripts in individual subjects BALF samples

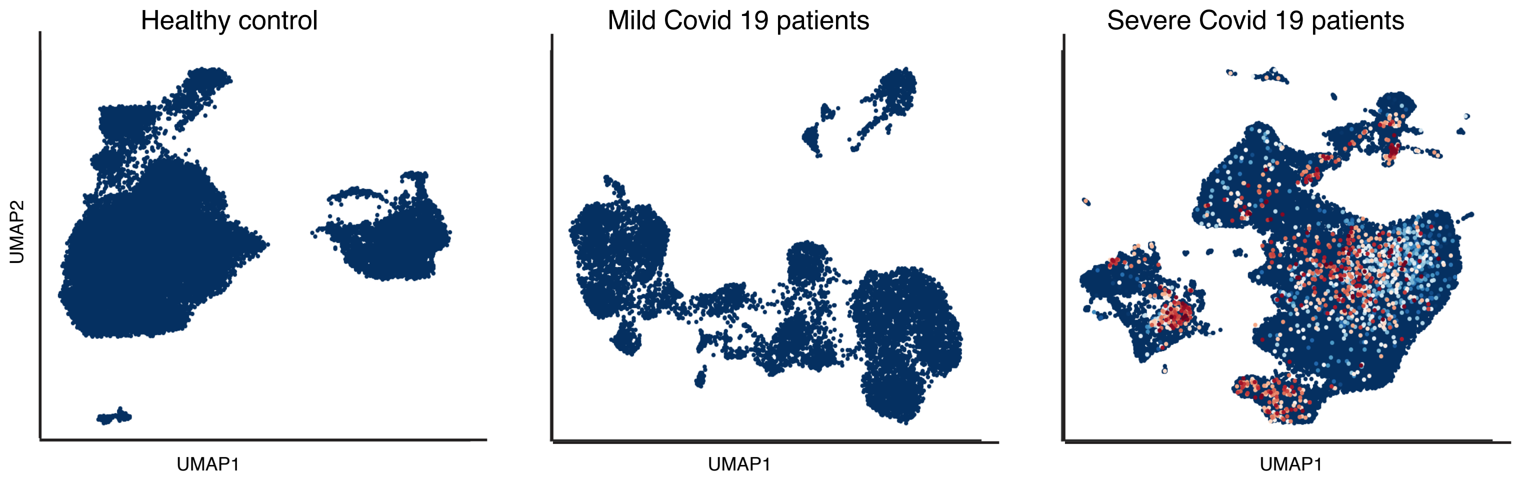


C. Target genes across subjects

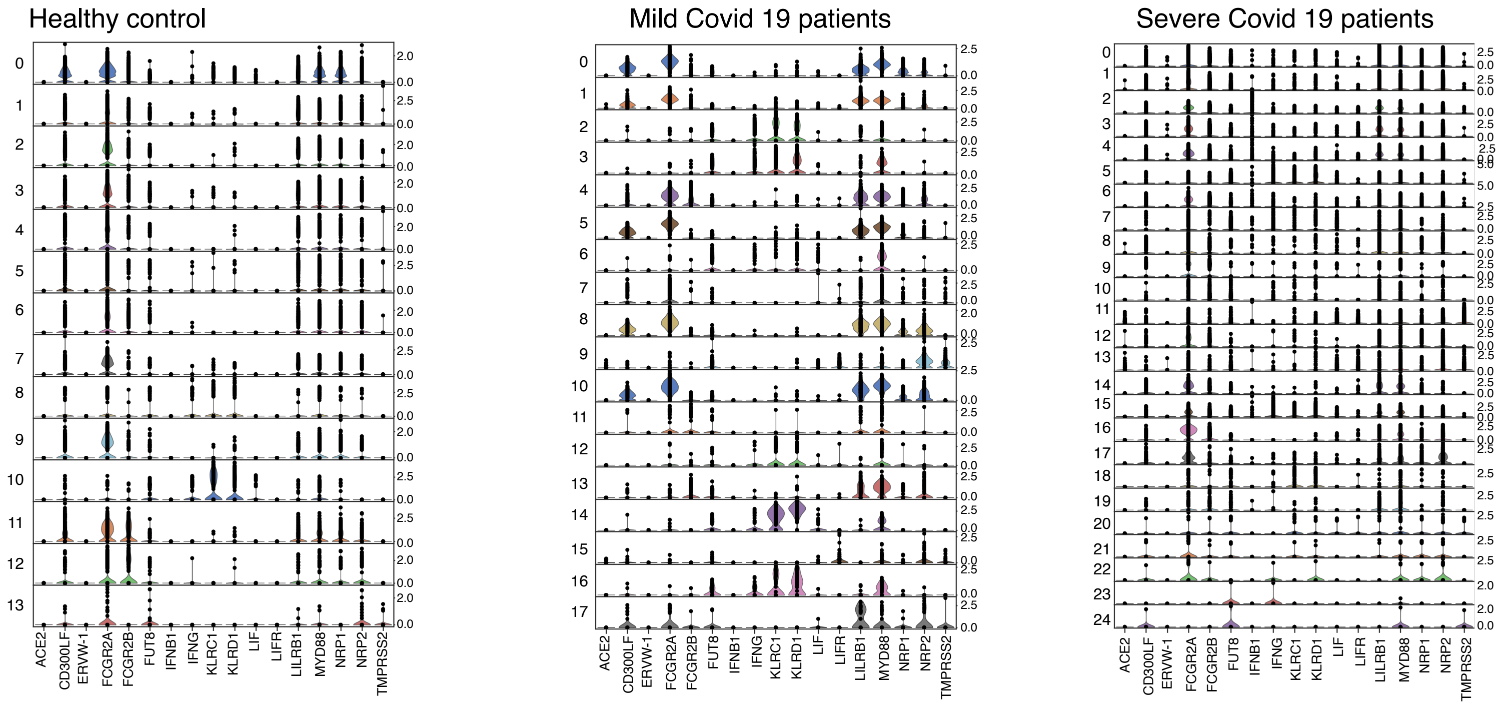
