## Supplementary Figure 7 for "Molecular relationships between SARS-CoV-2 Spike protein and LIFR, a pneumonia protective IL-6 family cytokine receptor"

### Figure S7 : Target genes expression in HCA lung derived single cells

A. Target genes expression in sub-sampled 300 individual cells across Alveolar Epithelial Type1 subpopulation (100 replications)

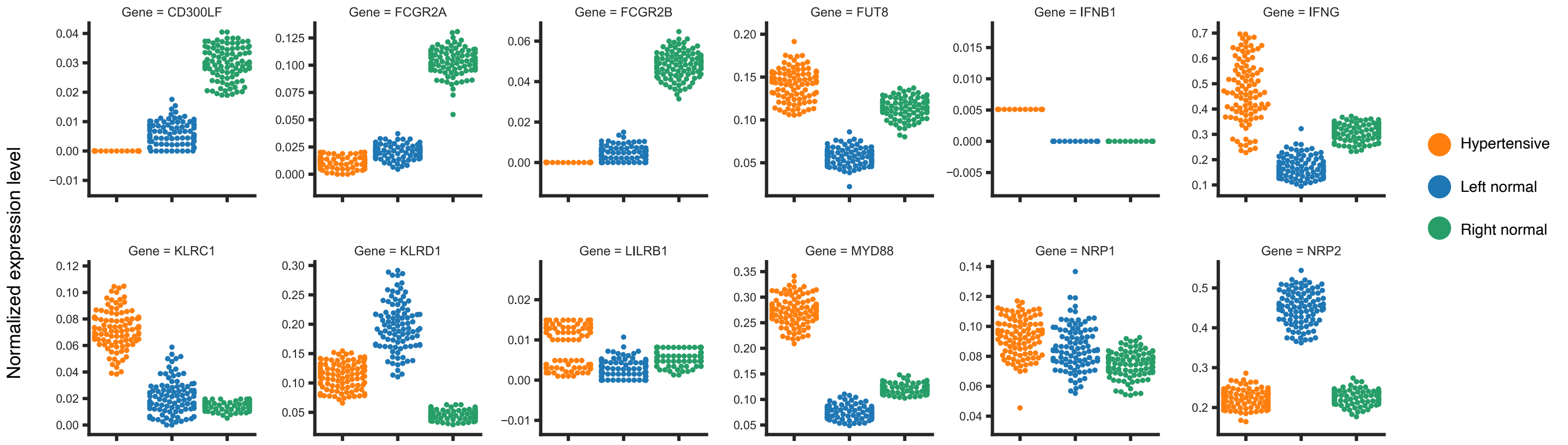


B. Target genes total counts in sub-sampled 300 individual cells across Alveolar Epithelial Type1 subpopulation (100 replications)

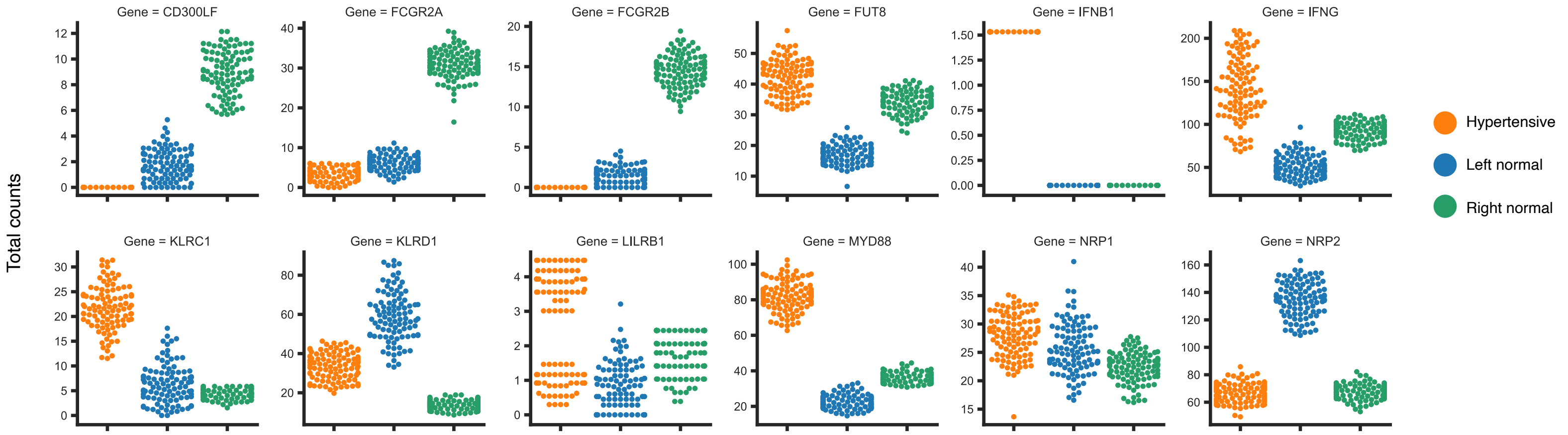


C. Target genes proportion of cells detected in sub-sampled 300 individual cells across Alveolar Epithelial Type1 subpopulation (100 replications)

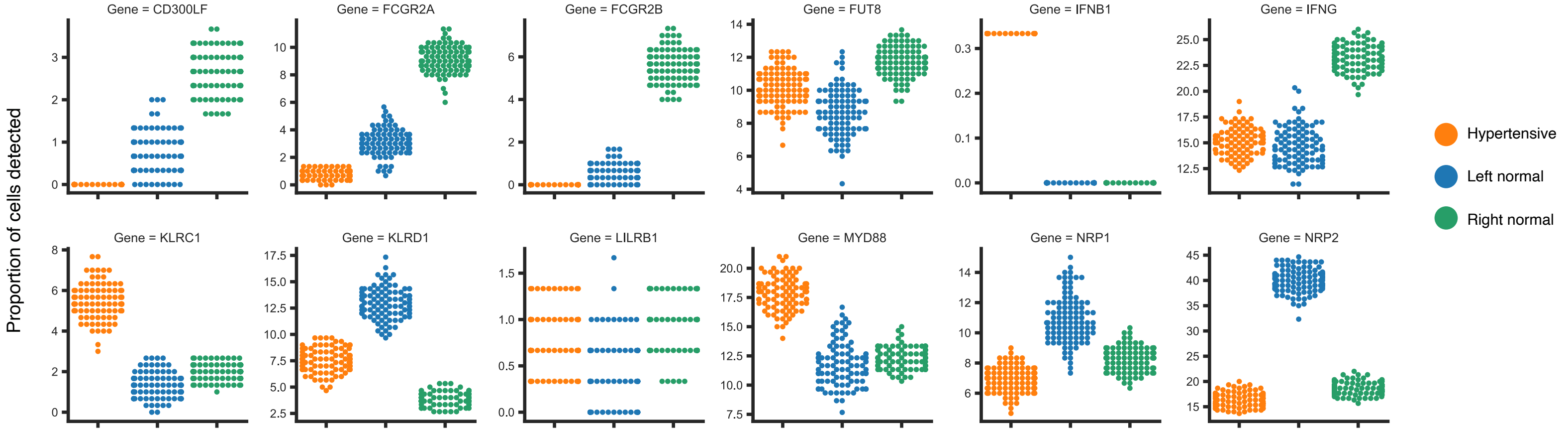
