## Supplementary Figure 8 for "Molecular relationships between SARS-CoV-2 Spike protein and LIFR, a pneumonia protective IL-6 family cytokine receptor"

### Figure S8 : Target genes expression in Covid19 and healthy controls at single cell resolution

#### Target genes across disease samples (Liao, M. et al., 2020 dataset)

A. Normalized mean expression in sub-sampled 300 cells across specific Ciliated Type1 subpopulation (100 replications)

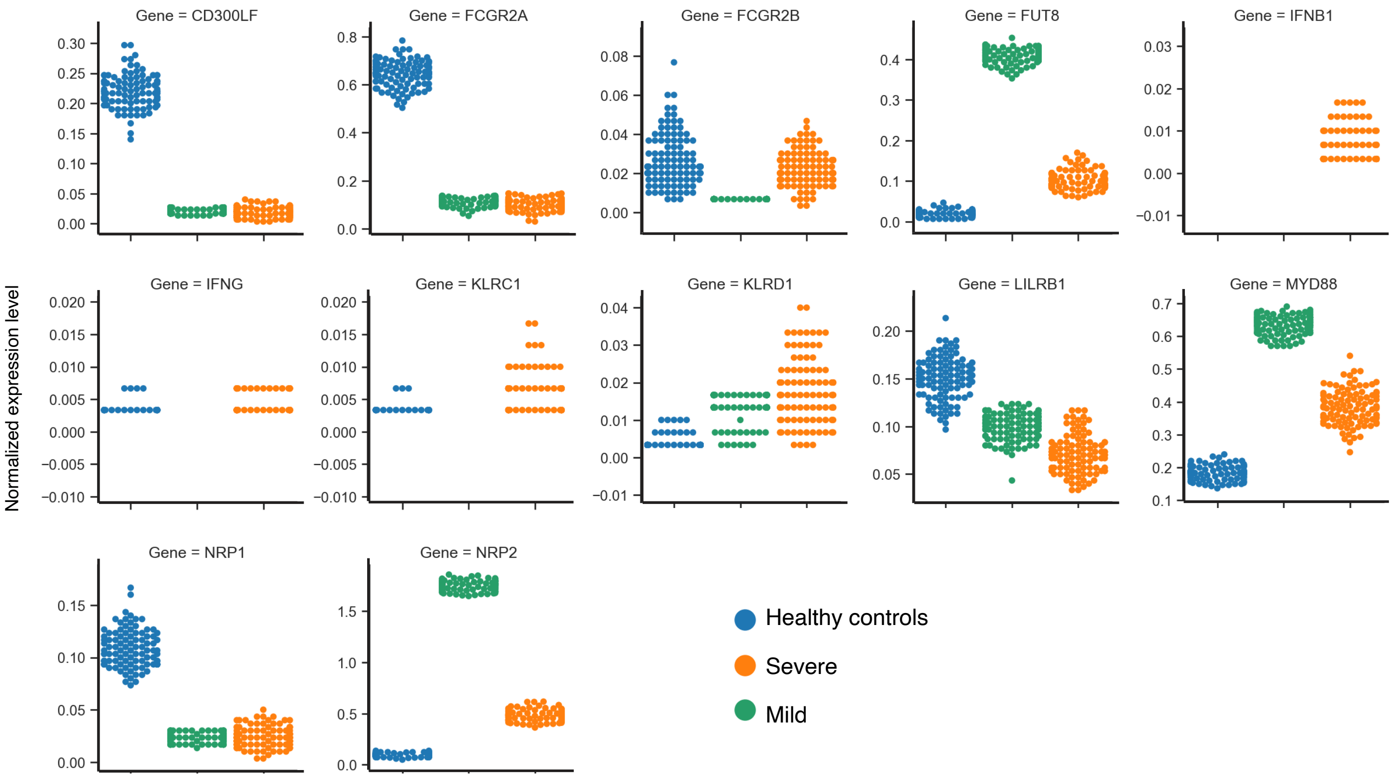
