## Supplementary Figure 9 for "Molecular relationships between SARS-CoV-2 Spike protein and LIFR, a pneumonia protective IL-6 family cytokine receptor"

### Figure S9 : Subpopulations cell type analysis in COVID19 datasets (Liao, M. et al., 2020 dataset)

#### A. Cell types in healthy controls

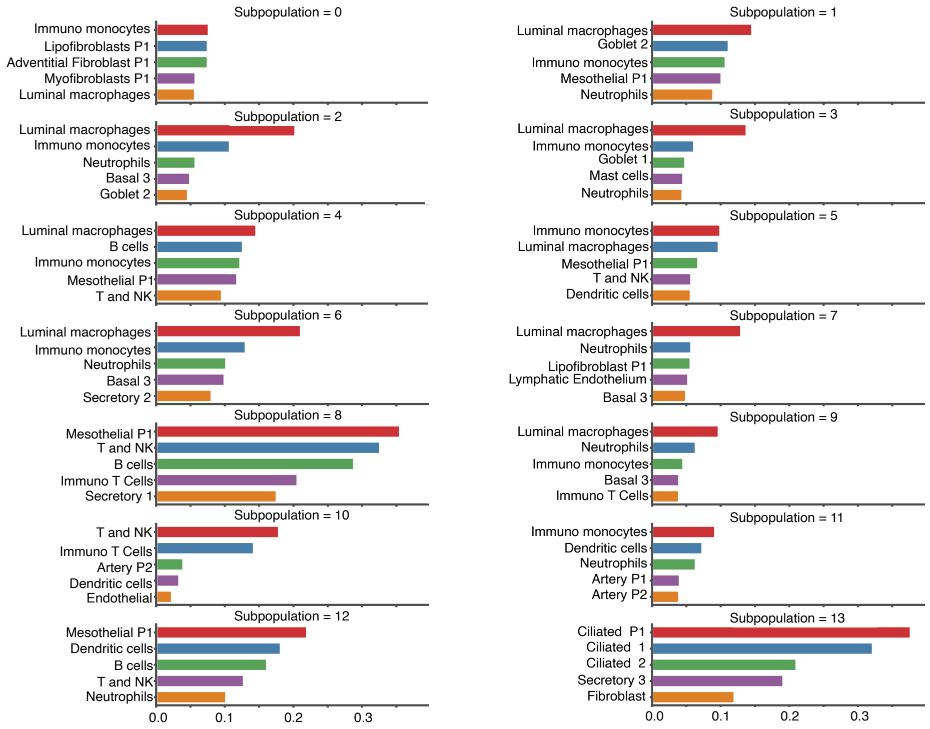


#### B. Cell types in mild COVID19 samples

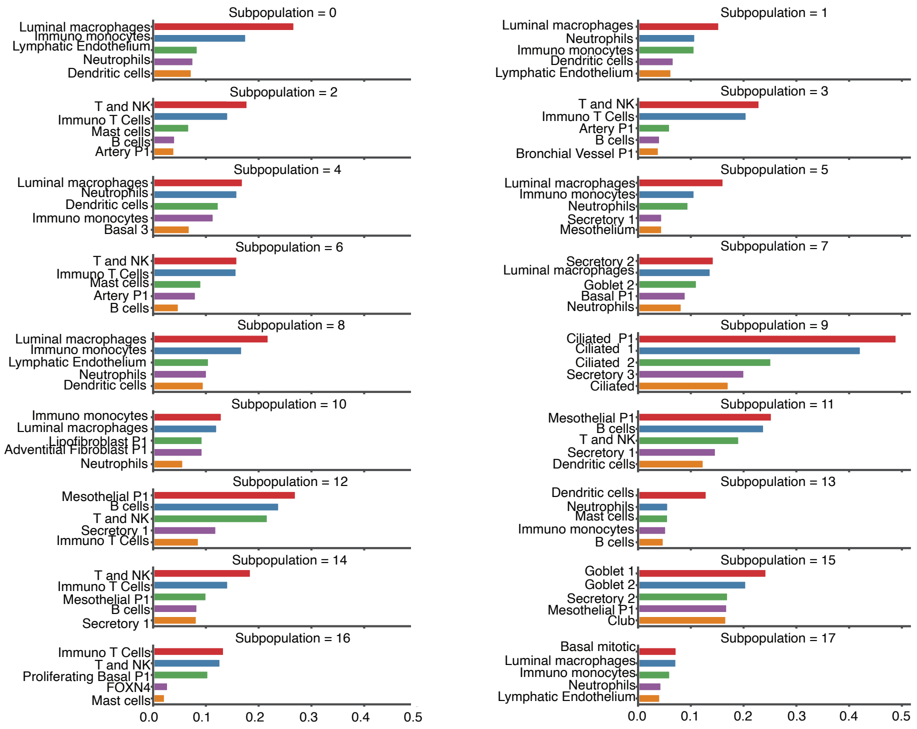


#### C. Cell types in severe COVID19 samples

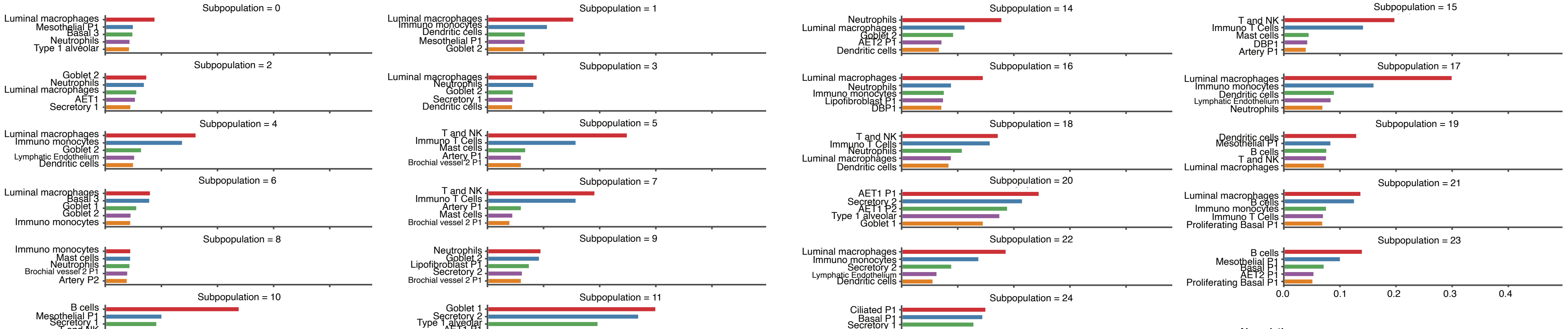


**Abreviation**:
DBP1 : Differentiating Basal P1
PBP1 : Proliferating Basal P1
AET1 P1 : Alveolar Epithelial Type1 P1
AET2 P1 : Alveolar Epithelial Type2 P1
